# Chromosomal-level genome assembly of the bioluminescent cardinalfish *Siphamia tubifer*, an emerging model for symbiosis research

**DOI:** 10.1101/2021.09.03.458932

**Authors:** AL Gould, JB Henderson, AW Lam

**Affiliations:** Ichthyology Department, Institute for Biodiversity Science and Sustainability, California Academy of Sciences, 55 Music Concourse Dr. San Francisco, CA 94118; Center for Comparative Genomics, Institute for Biodiversity Science and Sustainability, California Academy of Sciences, 55 Music Concourse Dr. San Francisco, CA 94118

## Abstract

The bioluminescent symbiosis between the sea urchin cardinalfish *Siphamia tubifer* (Kurtiformes: Apogonidae) and the luminous bacterium *Photobacterium mandapamensis* is an emerging vertebrate-bacteria model for the study of microbial symbiosis. However, there is little genetic data available for the host fish, limiting the scope of potential research that can be carried out with this association. In this study, we present a chromosomal-level genome assembly of *S. tubifer* using a combination of PacBio HiFi sequencing and Hi-C technologies. The final genome assembly was 1.2 Gb distributed on 23 chromosomes and contained 32,365 protein coding genes with a BUSCO completeness score of 99%. A comparison of the *S. tubifer* genome to that of another non-luminous cardinalfish revealed a high degree of synteny, whereas a similar comparison to a more distant relative in the Gobiiformes order revealed a fusion of two chromosomes in the cardinalfish genomes. An additional comparison of orthologous clusters among these three genomes revealed a set of 710 clusters that were unique to *S. tubifer* in which 23 GO pathways were significantly enriched, including several relating to host-microbe interactions and one involved in visceral muscle development, which could be related to the musculature involved in the gut-associated light organ of *S. tubifer*. We also assembled the complete mitogenome of *S. tubifer* and discovered both an inversion in the WANCY tRNA gene region resulting in a WACNY gene order as well as heteroplasmy in the length of the control region for this individual. A phylogenetic analysis based on the whole mitochondrial genome indicated that *S. tubifer* is divergent from the rest of the cardinalfish family, bringing up questions of the involvement of the bioluminescent symbiosis in the initial divergence of the ancestral *Siphamia* species. This draft genome assembly of *S. tubifer* will enable future studies investigating the evolution of bioluminescence in fishes as well as candidate genes involved in the symbiosis and will provide novel opportunities to use this system as a vertebrate-bacteria model for symbiosis research.

## Introduction

The cardinalfish genus *Siphamia* (Kurtiformes: Apogonidae) is comprised of 25 species, all of which are symbiotically bioluminescent. The fish has an abdominal light organ attached to the gut that harbors a dense population of a single species of luminous bacterium, *Photobacterium mandapamensis,* a member of the Vibrionaceae (Yoshiba & Haneda 1967, Wada *et al.* 2006, Kaeding *et al.* 2007, Urbanczyk *et al.* 2011, Gould *et al.* 2021). Additional cardinalfish species belonging to at least three other genera are also bioluminescent, however those species produce light autogenously and do not form a symbiosis with luminous bacteria (Thacker & Roje 2009). Members of the *Siphamia* genus are found throughout the Indo-Pacific, but *S. tubifer* (Figure 1) has the broadest distribution, spanning from east Africa to the French Polynesian Islands (Gon & Allen 2012). *Siphamia tubifer* is also the most well-studied *Siphamia* species to date; previous studies have characterized the fish’s life history (Gould *et al.* 2016), behavioral ecology (Eibl-Eibesfeldt 1961, Tamura 1982, Gould *et al.* 2014, 2015), and population genetics (Gould *et al.* 2017), as well as the the symbiosis with *P. mandapamensis* (Dunlap & Nakamura 2011, Dunlap *et al.* 2012, Gould *et al.* 2019, Iwai 1958, 1971). Unlike most symbiotically luminous fish species that inhabit deep water or have pelagic life histories, *S. tubifer* is a shallow, reef-dwelling species and can be raised in aquaria, both with and without its luminous symbiont, rendering the symbiosis to be experimentally tractable (Dunlap *et al.* 2012). Thus, the *S. tubifer-P. mandapamensis* symbiosis an emerging model for the study of vertebrate-bacteria associations, and is especially well-suited for studies of the vertebrate gut microbiome.

**Figure 1.**
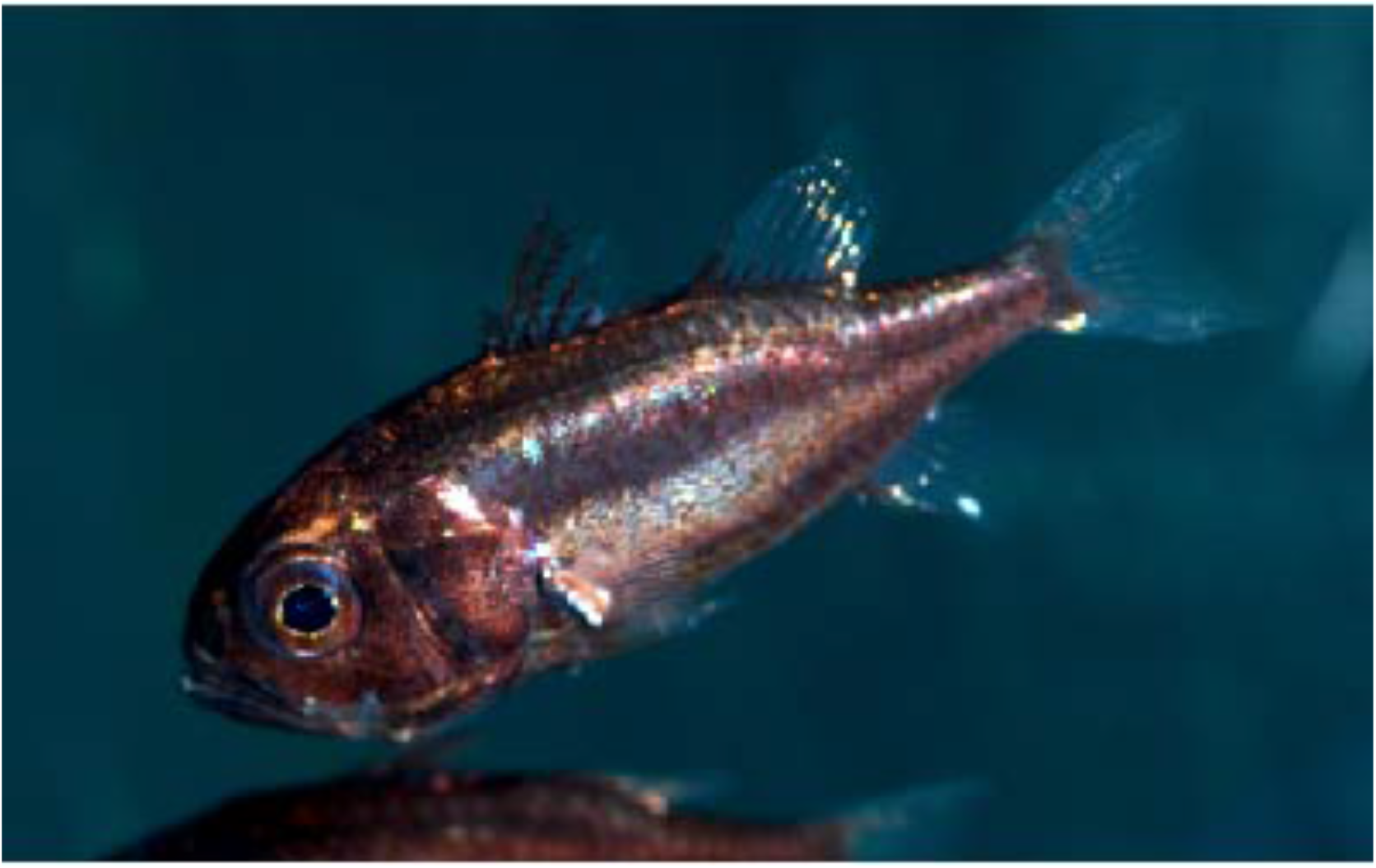
Photograph of *Siphamia tubifer.* Credit: Tim Wong, Steinhart Aquarium, California Academy of Sciences.

Despite an accumulation of knowledge of the biology of *S. tubifer* and its symbiosis with *P. mandapamensis*, there is little genomic information available for the fish, limiting the scope of possible studies that can be carried out with this association. A high-quality reference genome of *S. tubifer* will unlock new research opportunities to investigate the genetic mechanisms regulating this highly specific association, further enhancing its strength as a model system. Thus, we present a chromosomal-level assembly of the genome of *S. tubifer* produced by a combination of third-generation sequencing technology (PacBio HiFi sequencing) and chromosome conformation capture methods (Hi-C, Lieberman-Aide *et al.* 2009, vanBerkum *et al.* 2010). We then compare our *S. tubifer* genome assembly and annotation to that of other chromosomal-level assemblies of a closely related but non-luminous cardinalfish species and a more distant relative in the sister order Gobiiformes to describe synteny between the genomes and identify candidate genes that could be involved in the symbiosis. We also present a whole mitochondrial genome assembly of *S. tubifer* and use this sequence information to infer *S. tubifer*’s phylogenetic position within the cardinalfish family, providing further insight into the evolution of this bioluminescent symbiosis.

## Methods

### Tissue collection, DNA extraction and sequencing

All tissue was obtained from a single female *Siphamia tubifer* specimen collected from a shallow reef in Okinawa, Japan (26.66°N, 127.88°E). The fish was collected and euthanized following approved protocols and permits for the capture, care and handling of fish by the California Academy of Science’s Institutional Animal Care and Use Committee. Immediately following euthanasia, fresh muscle tissue was sampled from the flanking region of the fish for high molecular weight (HMW) DNA extraction using a phenol-chloroform extraction protocol provided by Pacific Biosciences of California, Inc. Fresh muscle and brain tissue were also sampled from the same individual for Hi-C methods. The HMW DNA was prepared for PacBio HiFi sequencing at UC Berkeley’s QB3 Genomics Sequencing Lab (Berkeley, CA) and sequenced on one Sequel II 8M SMRT Cell.

### Hi-C library preparation and sequencing

*In situ* Hi-C libraries were prepared from the freshly homogenized muscle and brain tissues following the protocol described in Rao *et al.* (2014) with slight modifications. After the Streptavidin pull-down step, the biotinylated Hi-C products underwent end repair, ligation, and enrichment using the NEBNext^®^ Ultra™II DNA Library Preparation kit (New England Biolabs Inc, Ipswich, MA). Titration of the number of PCR cycles was performed as described in Belton *et al.* (2012). The final libraries were then sequenced as paired-end 150 bp reads on the Illumina NovaSeq 6000 platform by Novogene Corporation, Inc. (Sacramento, CA).

### Genome size estimation, assembly and chromosome mapping

Circular Consensus Sequences (CCS) were generated using ccs v5.0.0 (https://github.com/PacificBiosciences/pbbioconda), from 35.95M subreads, representing 442.25G bases, and filtered to produce HiFi reads, defined as having at least two circular passes and minimum of 99.9% accuracy. A custom script created a .fastq file containing the HiFi reads extracted from the .bam output file of the ccs step. Jellyfish (Marcais & Kingsford 2012) was then used to count and create histograms of kmers size 21 and 25 from the HiFi reads, and GenomeScope v2.0 (Ranallo-Benavidez *et al.* 2020) was run on each set to determine estimates of genome size.

Next, filtering was performed to remove contaminant sequences. Since using blastn (Altschul *et al.* 1990) and other similar tools is inefficient with long reads, we first used minimap2 (Li 2018) with the genome of the closely related orbiculate cardinalfish, *Sphaeramia orbicularis*, to exclude matching reads from further contaminant analysis. For the remaining sequences, blastn was then used against a database of *Siphamia tubifer*’s luminous symbiont, *Photobacterium mandapamensis* (Urbanczyk *et al.* 2011), to identify its sequences as contaminants. Additionally, to further reduce the analysis, the first 500 bases of the remaining long reads were used as blastn queries against the nt database with option -taxidlist restricting search to bacteria, and those excluded with e-value greater than −1e10. Similarly, mitochondrial DNA sequences were identified and removed for separate analysis by using blastn against a database of three Apogonidae mitochondrial genomes: *S. orbicularis*, *Ostorhinchus fleurieu*, and *Pristicon trimaculatus*. Subsequent nuclear genome analysis used the remaining long read HiFi sequences with contaminant and mitochondrial sequences removed.

The remaining HiFi sequences were assembled with hifiasm v0.13-r308 (Cheng *et al.* 2021). The hifiasm assembly program is designed for HiFi reads produced from a diploid genome and also incorporates purge_dups (Guan *et al.* 2020) to separate out duplicate haplotigs, producing a primary assembly of the higher quality contigs and an alternate assembly of contigs including the duplicates. For comparison, we also ran Improved Phase Assembler, ipa v1.3.0, (PacificBiosciences 2020) to create an assembly from the same input. We then ran quickmerge v0.3 (Chakraborty *et al.* 2016) for a third assembly where the hifiasm result was used as the query and the ipa output as a reference assembly to attempt to bridge gaps in the hifiasm genome representation.

The Hi-C reads, consisting of 624.35M combined brain and muscle tissue read pairs, were mapped using juicer v1.6 (Durand *et al*. 2016b) against the hifiasm assembled contig level genome. We next ran 3d-dna v180922 (Dudchenko *et al.* 2017) with its early-exit flag to create an input file for JuiceBox Assembly Tools (JBAT) (Durand *et al.* 2016a, Dudchenko *et al*. 2018) that represents the assembly with contigs ordered and oriented in a candidate chromosomal level depiction. Using JBAT, we interactively updated location and orientation of contigs and their delineation at the chromosome level (Figure 2a). This assembly was also queried against the nt database using blastn to identify any additional contaminants for removal.

**Figure 2.**
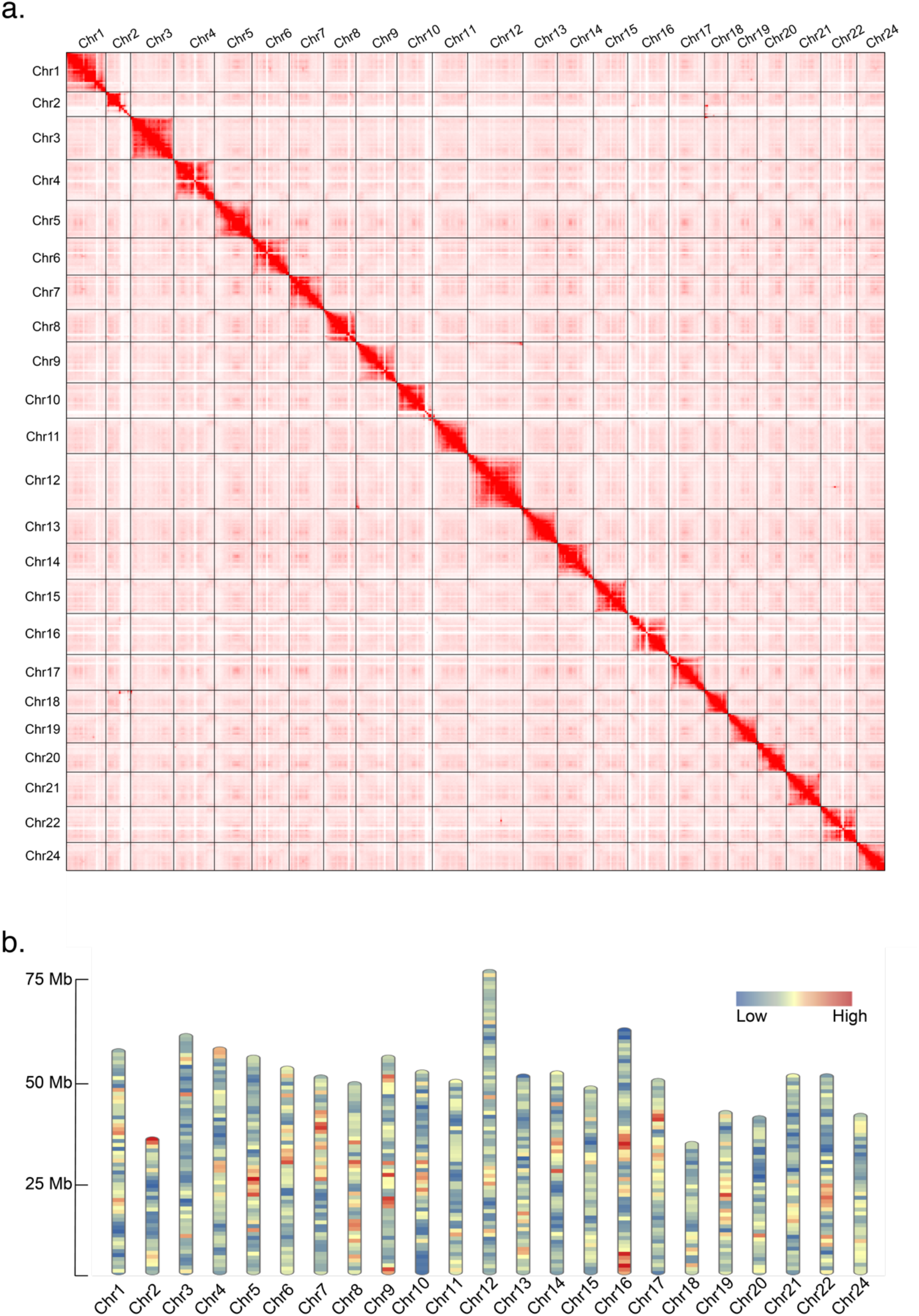
**a)** Hi-C contact heatmap for *Siphamia tubifer*. Black lines indicate chromosome boundaries. **b)** Gene density distributed across the 23 chromosomes of the *S. tubifer* genome.

To assess the level of genome completeness, we ran BUSCO v5.12 (Simão *et al.* 2015) with the 3,640 entry Actinopterygii dataset in both its MetaEuk (Karin *et al.* 2020) and AUGUSTUS (Keller *et al.* 2011) modes. We then used a custom script to update BUSCOs found by AUGUSTUS that were missing in the MetaEuk results and another to report the combined BUSCO scores.

### Gene annotation and synteny

Prior to gene annotation, *de novo* repeats were identified from the *S. tubifer* genome assembly using RepeatModeler v2.0.1(Flynn *et al.* 2020). First, the .fasta file representing these species specific repeat models and the vertebrate repeat models from Repbase (https://www.repeatmasker.org) RepeatMasker libraries v20181026 were appended into a combined file. This file was then used as the input library to Repeatmasker v4.0.9 (Smit *et al.* 2013-2015) with the options -small -xsmall and -nolow to create the soft-masked repeat version of the assembly file used for gene model annotation. BRAKER2 (Brůna *et al.* 2021), using GeneMark-EP+ (Brůna *et al.* 2020) and AUGUSTUS, combined with the vertebrate protein database from OrthoDB v10 (https://www.orthodb.org) (Kriventseva *et al.* 2019), was used for gene annotation. The output of potential gene models represented in .gff3, amino acid, and DNA files were subject to additional filtering together with functional annotation.

To check for protein domains, we ran InterProScan v5.51-85.0 (Jones *et al.* 2014) on the amino acid sequences found in the BRAKER2 results. These sequences were also used as queries for a blastp run on three databases: SwissProt, TrEMBL, and the vertebrate proteins from OrthoDB v10. The DNA versions of the sequences were also queried with blastn against the nt database downloaded on February 13, 2021. Gene models, in .gff3, amino acid, and DNA files, were kept for those sequences with an InterProScan determined protein domain and one of the four database searches yielding a match with an e-value 0.1e-6 or less. These files were then updated with the matching descriptions indicating they were similar to the highest scoring match of the four searches. Protein domain IDs and Gene Ontology (GO) terms, as determined by the InterProScan output, were added to the .gff3 file for each retained gene model as was the functional annotation description. tRNAscan-SE v2.0.8 (Chan *et al.* 2021) was implemented to identify tRNAs throughout the genome.

We then compared the genome annotation of *S. tubifer* to that of the closely related non-luminous cardinalfish, *Sphaeramia orbicularis*, and to a more distant member of the sister order Gobiiformes, the mudskipper *Periophthalmus magnuspinnatus,* using OrthoVenn2 (Xu *et al.* 2019). We determined the number of shared and unique protein clusters among these species and carried out a GO enrichment analysis on the unique clusters identified for *S. tubifer.* Next, we examined synteny between our *S. tubifer* genome assembly and the chromosomal-level genomes of both *S. orbicularis* (GenBank GCF_902148855.1) *and P. magnuspinnatus* (GenBank GCA_009829125.1) using the set of single copy orthologs identified from the BUSCO (Simão *et al.* 2015) Actinopterygii gene set and converted the output for visualization in Circos (Krzywinski *et al.* 2009) using custom scripts.

### Mitochondrial genome assembly and analysis

Mitochondrial genome analysis was based on sequences matching at least 60% query coverage in a blastn match (qcovus format specifier) to one of the three Apogonidae mitochondrial genomes; *S. orbicularis*, *O. fleurieu*, and *P. trimaculatus*. When matched to the reverse strand, sequences were reverse complimented and _RC was appended to the name, resulting in all sequences having the same strand orientation. Megahit (Li *et al.* 2015) was then run on these sequences to assemble a draft mitogenome and MITOS2 (Bernt *et al.* 2013) was used to annotate the mitogenome.

GenBank annotations for the three Apogonidae mitogenomes were downloaded and their sequences were extracted into .fasta files containing records corresponding to the genome’s rRNAs, tRNAs, and protein coding genes. The *S. tubifer* mitochondrial HiFi reads were queried with a subject database of these sequences from the three mitogenomes using blastn with its - task blastn option (overriding default -task megablast). These matches were then used to split reads into three sets of new .fasta records using tRNA *Phe* and *Pro* as markers: (a) *Phe* to *Pro* (or end of read if no *Pro*), (b) if no *Phe*, then beginning of read to *Pro*, (c) Pro to Phe when both found, capturing the complete control region in between. The first 2 sets were used for tRNA analysis, including tRNA order, and the third set was used for control region repeat and heteroplasmy analysis. Mitfi (Jühling *et al.* 2012) was used to identify tRNAs from 176 reads from sets (a) and (b) that matched at least 90% query coverage to one of the three closely related species’ mitogenomes. Tandem Repeat Finder (TRF) (Bensen 1999) was run to find repeats in the control region set.

Using the whole mitochondrial genome assembly, the phylogenetic placement of *S. tubifer* within the cardinalfish family was inferred. This analysis also included one *Kurtus* species and several species of gobies for reference, as well as two members of the Syngnathiformes order as an outgroup. Whole mitochondrial sequences (excluding the control regions) were aligned using MAFFT (Katoh *et al.* 2002), and the aligned reads were used to construct maximum likelihood trees with raxml-ng (Kozlov *et al.* 2019) using the substitution model with the lowest BIC score as predicted by IQtree (Nguyen *et al* 2015) and 500 bootstrap replicates.

## Results

### Genome size estimation, assembly, and chromosome mapping

A total of 2,110,443 HiFi CCS reads consisting of 27,799,385,228 bp were generated from the single HiFi library, with a polymerase N50 of 183,061 and subread N50 of 13,439. Over 97% of the HiFi reads were between 12,000 and 15,000 bp. From these sequences, the GenomeScope size estimate, using kmer lengths 21 and 25, ranged from 947,587,691 to 964,260,239 bp. Repeat length was estimated as 215,783,447 to 256,391,534 bp, though repeat length is often underestimated by kmer counting models, leading to a lower overall estimate of genome size. After contaminant and mitochondrial sequence removal 2,109,973 sequence reads were left with 6,158,291 bp excluded from the source HiFi reads. These remaining sequences were used as input for the assembly programs hifiasm and ipa. Based on contiguity and accuracy metrics, we used the hifiasm assembly to scaffold with the Hi-C reads.

For the Hi-C libraries, a total of 742,280,226 and 506,411,380 reads were produced from the muscle and brain tissue, respectively. Of those, 100% of the muscle reads and 99.98% of the brain reads were clean and of high quality, resulting in GC contents of 39.3% and 43.9%. The Juicer mapping program found 245,145,667 read pairs having Hi-C contacts. After interactive modification with JBAT, guided by the 3d-dna program contig placement and orientation, the resulting genome assembly was 1.2 Gb distributed on 23 chromosomes, and 1.81% unplaced scaffolds, with a contig N50 of 2.3 Mb and scaffold N50 of 51.1 Mb (Table 1), and 37.71% GC content. There are 1,960 contigs constituting chromosomal sequences. An additional two dozen smaller contig records were identified as contaminants by the final nt blastn search (primarily Arthropoda, though of unknown origin) and were removed to produce the final assembly. This assembly has the same summary statistics reported above except for the unplaced scaffold percentage, which was slightly lower (1.74%).

**Table 1.** Assembly statistics of the draft genome of *Siphamia tubifer*.

|  |  |
| --- | --- |
| **Assembly size** | 1,200,827,456 bp |
| **Total scaffolds** | 118 |
| **Total contigs** | 2,057 |
| **Scaffold N50 (L50)** | 51,162,488 bp (11) |
| **Contig N50 (L50)** | 2,340,513 bp (133) |
| **Scaffold N90 (L90)** | 40,504,042 bp (21) |
| **Contig N90 (L90)** | 318,326 bp (689) |
| **Total genes** | 32,364 |
| **Mean gene length** | 12,504 bp |

The 23 chromosomes in the *S. tubifer* genome assembly are numbered 1 to 22 and 24 based on synteny with another cardinalfish genome, the 23 chromosome *S. orbicularis* genome assembly fSphaOr1.1 (GenBank GCF_902148855.1), which is based on synteny with the 24 chromosome medaka genome (GenBank ASM223467v1), representing the fusion of the medaka chr23 into a cardinalfish chromosome.

BUSCO completeness assessment from the 3,640 entry Actinopterygii dataset show 99% complete with just 13 of the genes not found (MetaEuk mode: 98% complete, AUGUSTUS mode: 97.2% complete).

### Genome annotation and statistics

Repeat analysis indicated 626,216,533 bp, or 52.11% of the genome, classified as repeats, of which, most (23.7% of the genome) are DNA repeat elements. Additionally, 7.03% of the genome contains long interspersed nuclear elements (LINEs), with 16.28% of the genome characterized as unclassified repeats. The extent of repeats may account for the discrepancy between the assembly size and the GenomeScope estimates using kmer counts.

Gene annotation identified 30,117 gene models with a total length of 360,171,123 bp, (29.99% of the genome). Exons at 53,076,342 bp are 4.42% of the genome and average 9.64 per gene; fewer than 10% are single exon genes. Additional per chromosome details of genes, exons, and introns are outlined in Table 2. The orbiculate cardinalfish, *S. orbicularis* (GenBank Annotation Release 100 2019-08-03), was the closest functional annotation reference for 17,079 (56.7%) of the 30,117 *S. tubifer* gene models. This was followed by several other fish species: *Lates calcarifer* (n=2,317), *Seriola dumerili* (n=1,357), *Larimichthys crocea* (n=995), and *Stegastes partitus* (n=779).

**Table 2.**
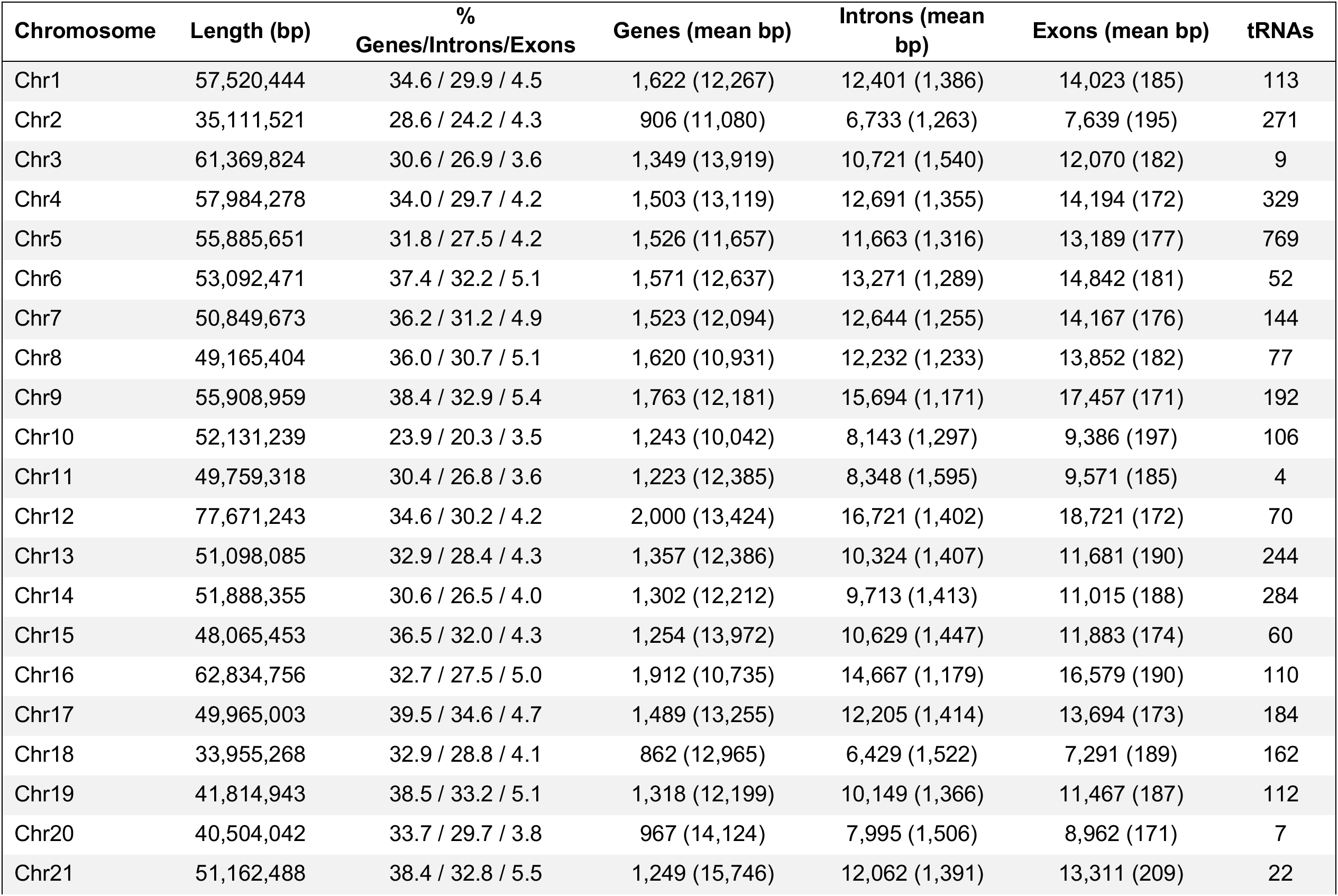
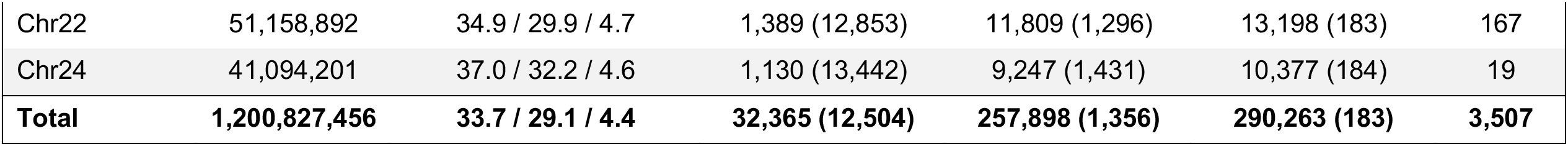
Annotation statistics of the *S. tubifer* genome by chromosome. For each chromosome the total length in bp and the percent of those bp belonging to genes, introns, and exons are listed as well as the number of genes, introns, exons and tRNAs. The mean length of genes, introns, and exons in each chromosome are also included.

| Chromosome | Length (bp) | %<br>Genes/Introns/Exons | Genes (mean bp) | Introns (mean<br>bp) | Exons (mean bp) | tRNAs |
| --- | --- | --- | --- | --- | --- | --- |
| Chr1 | 57,520,444 | 34.6 / 29.9 / 4.5 | 1,622 (12,267) | 12,401 (1,386) | 14,023 (185) | 113 |
| Chr2 | 35,111,521 | 28.6 / 24.2 / 4.3 | 906 (11,080) | 6,733 (1,263) | 7,639 (195) | 271 |
| Chr3 | 61,369,824 | 30.6 / 26.9 / 3.6 | 1,349 (13,919) | 10,721 (1,540) | 12,070 (182) | 9 |
| Chr4 | 57,984,278 | 34.0 / 29.7 / 4.2 | 1,503 (13,119) | 12,691 (1,355) | 14,194 (172) | 329 |
| Chr5 | 55,885,651 | 31.8 / 27.5 / 4.2 | 1,526 (11,657) | 11,663 (1,316) | 13,189 (177) | 769 |
| Chr6 | 53,092,471 | 37.4 / 32.2 / 5.1 | 1,571 (12,637) | 13,271 (1,289) | 14,842 (181) | 52 |
| Chr7 | 50,849,673 | 36.2 / 31.2 / 4.9 | 1,523 (12,094) | 12,644 (1,255) | 14,167 (176) | 144 |
| Chr8 | 49,165,404 | 36.0 / 30.7 / 5.1 | 1,620 (10,931) | 12,232 (1,233) | 13,852 (182) | 77 |
| Chr9 | 55,908,959 | 38.4 / 32.9 / 5.4 | 1,763 (12,181) | 15,694 (1,171) | 17,457 (171) | 192 |
| Chr10 | 52,131,239 | 23.9 / 20.3 / 3.5 | 1,243 (10,042) | 8,143 (1,297) | 9,386 (197) | 106 |
| Chr11 | 49,759,318 | 30.4 / 26.8 / 3.6 | 1,223 (12,385) | 8,348 (1,595) | 9,571 (185) | 4 |
| Chr12 | 77,671,243 | 34.6 / 30.2 / 4.2 | 2,000 (13,424) | 16,721 (1,402) | 18,721 (172) | 70 |
| Chr13 | 51,098,085 | 32.9 / 28.4 / 4.3 | 1,357 (12,386) | 10,324 (1,407) | 11,681 (190) | 244 |
| Chr14 | 51,888,355 | 30.6 / 26.5 / 4.0 | 1,302 (12,212) | 9,713 (1,413) | 11,015 (188) | 284 |
| Chr15 | 48,065,453 | 36.5 / 32.0 / 4.3 | 1,254 (13,972) | 10,629 (1,447) | 11,883 (174) | 60 |
| Chr16 | 62,834,756 | 32.7 / 27.5 / 5.0 | 1,912 (10,735) | 14,667 (1,179) | 16,579 (190) | 110 |
| Chr17 | 49,965,003 | 39.5 / 34.6 / 4.7 | 1,489 (13,255) | 12,205 (1,414) | 13,694 (173) | 184 |
| Chr18 | 33,955,268 | 32.9 / 28.8 / 4.1 | 862 (12,965) | 6,429 (1,522) | 7,291 (189) | 162 |
| Chr19 | 41,814,943 | 38.5 / 33.2 / 5.1 | 1,318 (12,199) | 10,149 (1,366) | 11,467 (187) | 112 |
| Chr20 | 40,504,042 | 33.7 / 29.7 / 3.8 | 967 (14,124) | 7,995 (1,506) | 8,962 (171) | 7 |
| Chr21 | 51,162,488 | 38.4 / 32.8 / 5.5 | 1,249 (15,746) | 12,062 (1,391) | 13,311 (209) | 22 |
| Chr22 | 51,158,892 | 34.9 / 29.9 / 4.7 | 1,389 (12,853) | 11,809 (1,296) | 13,198 (183) | 167 |
| Chr24 | 41,094,201 | 37.0 / 32.2 / 4.6 | 1,130 (13,442) | 9,247 (1,431) | 10,377 (184) | 19 |
| **Total** | **1,200,827,456** | **33.7 / 29.1 / 4.4** | **32,365 (12,504)** | **257,898 (1,356)** | **290,263 (183)** | **3,507** |

The orthologous cluster analysis indicated that a much lower number of protein clusters were shared between *S. tubifer* and the mudskipper *P. magnuspinnatus* (n=419) than with the other cardinalfish *S. orbicularis* (n=1,743). However, *S. orbicularis* shared a much larger number of clusters with *P. magnuspinnatus* (n=1,484) than did *S. tubifer*. There were also 710 unique protein clusters that were present only in the *S. tubifer* genome (Figure 3a), of which 506 were assigned to GO categories (Table S1). Overall, the largest percent of these clusters were categorized as biological processes (GO:0008150) (26%) and cellular processes (GO:0009987) (16%), and another 8% and 5% were identified as response to stimulus (GO:0050896) and developmental processes (GO:0032502), respectively (Figure 3b). The largest cluster was made up of 70 proteins assigned as DNA integration (GO:0015074), and the second largest cluster contained 41 proteins relating to visual perception (GO:0007601). There were also 9 genes unique to *S. tubifer* that were categorized as immune system processes (GO:0002376) (Table S2). An enrichment analysis of these unique clusters also revealed 26 functions that were significantly enriched (p>0.01). Of those, several were related to viral penetration (GO:0075732) and integration (GO:0044826) into a host as well as visceral muscle development (GO:0007522) (Table 3).

**Figure 3.**
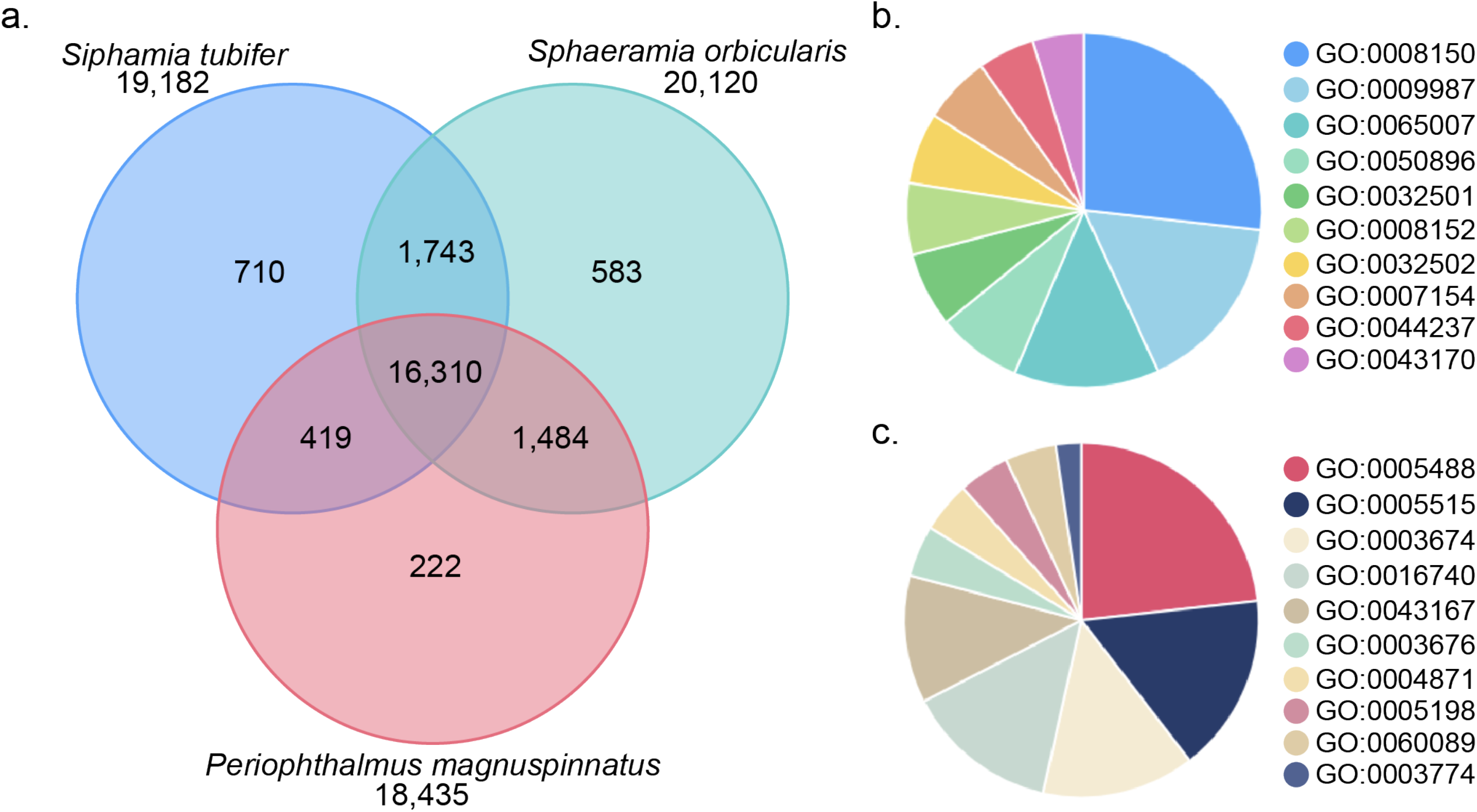
**a)** Venn diagram of the distribution of orthologous clusters among *Siphamia tubifer*, the non-luminous cardinalfish *Sphaeramia orbicularis*, and the mudskipper *Periophthalmus magnuspinnatus* (order Gobiiformes). **b)** Distribution of the top ten biological process GO terms assigned to the 710 unique clusters identified for *S. tubifer* and **c)** the top ten molecular function GO terms assigned to the gene clusters

**Table 3.** Gene ontology (GO) enrichment analysis for the 710 unique clusters identified in the *Siphamia tubifer* genome from a three-way comparison of orthologous clusters with the non-luminous cardinalfish *Sphaeramia orbicularis* and the mudskipper *Periophthalmus magnuspinnatus* (order Gobiiformes).

| GO ID | Description | Count | p-value |
| --- | --- | --- | --- |
| GO:0071625 | vocalization behavior | 6 | 7.13E-07 |
| GO:0050808 | synapse organization | 8 | 8.31E-07 |
| GO:0006310 | DNA recombination | 5 | 2.52E-06 |
| GO:0015074 | DNA integration | 4 | 6.32E-06 |
| GO:0030050 | vesicle transport along actin filament | 4 | 2.80E-05 |
| GO:0006313 | transposition, DNA-mediated | 4 | 4.91E-05 |
| GO:0006953 | acute-phase response | 3 | 5.44E-04 |
| GO:0075732 | viral penetration into host nucleus | 2 | 6.69E-04 |
| GO:0032185 | septin cytoskeleton organization | 2 | 6.69E-04 |
| GO:0001764 | neuron migration | 3 | 1.24E-03 |
| GO:0071805 | potassium ion transmembrane transport | 3 | 1.24E-03 |
| GO:0034765 | regulation of ion transmembrane transport | 8 | 1.35E-03 |
| GO:0003964 | RNA-directed DNA polymerase activity | 2 | 1.95E-03 |
| GO:0043516 | regulation of DNA damage response, signal transduction by p53 class mediator | 2 | 1.95E-03 |
| GO:0044826 | viral genome integration into host DNA | 2 | 1.95E-03 |
| GO:1900044 | regulation of protein K63-linked ubiquitination | 2 | 1.95E-03 |
| GO:0032197 | transposition, RNA-mediated | 2 | 1.95E-03 |
| GO:0047369 | succinate-hydroxymethylglutarate CoA-transferase activity | 2 | 1.95E-03 |
| GO:0007522 | visceral muscle development | 2 | 1.95E-03 |
| GO:0006261 | DNA-dependent DNA replication | 2 | 1.95E-03 |
| GO:0090129 | positive regulation of synapse maturation | 2 | 1.95E-03 |
| GO:0043551 | regulation of phosphatidylinositol 3-kinase activity | 2 | 1.95E-03 |
| GO:0045162 | clustering of voltage-gated sodium channels | 2 | 1.95E-03 |
| GO:0045214 | sarcomere organization | 4 | 2.03E-03 |
| GO:0015031 | protein transport | 10 | 3.13E-03 |
| GO:0008154 | actin polymerization or depolymerization | 2 | 3.81E-03 |

### Genome synteny

Overall a high degree of synteny between the genomes of *S. tubifer* and the nonluminous, orbiculate cardinalfish *S. orbicularis* was observed (Figure 4a). Of the 3,555 orthologous genes from the BUSCO set only 2.5% (n=90) changed chromosomal assignment. A comparison to a more distantly related fish species, the mudskipper *P. magnuspinnatus,* a member of the sister order Gobiiformes revealed that a merge occurred between *P. magnuspinnatus* chromosomes 12 and 23 to become chromosome 12 in both cardinalfish genomes (Figure 4b). Thus, the mudskipper genome has one more chromosome (n=24) than both *S. tubifer* and *S. orbicularis* (n=23).

**Figure 4.**
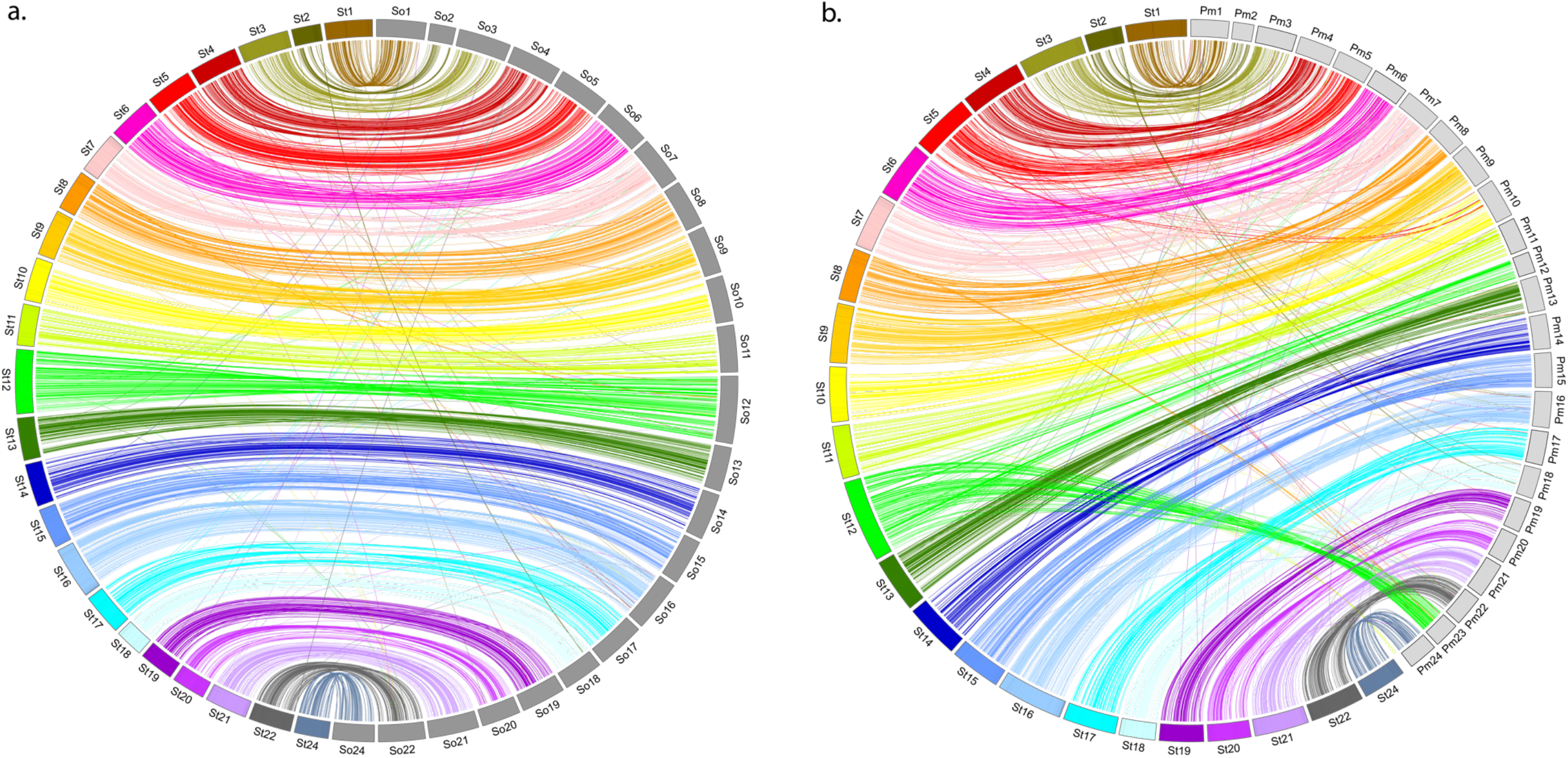
Circos plots depicting synteny between the genomes of *Siphamia tubifer* and **a)** the orbiculate cardinalfish, *Sphaeramia orbicularis* and **b)** the mudskipper *Periophthalmus magnuspinnatus*. Each chromosome in the *S. tubifer* genome is represented by a distinct color whereas the *S. orbicularis* and *P. magnuspinnatus* chromosomes are shown in dark and light gray, respectively. Links between single copy orthologs from the BUSCO Actinopterygii gene set are shown.

### Mitochondrial genome

There were 5,124,329 total bp in the 392 HiFi reads that matched the cut-off of 60% query coverage used in the mitochondrial sequence analysis. Assuming a mitogenome is between 16,000 and 18,000 bp, this represents 285-320x coverage. There were 176 reads in which 90% or greater of the read length was covered containing 2,302,235 bp.

The complete mitochondrial genome averaged 17,905 bp, but varied due to heteroplasmy in the length of the control region (Figure 5a). There were 13 protein coding genes, 22 tRNA genes, and 2 rRNA genes, as expected for a vertebrate mitogenome. However, there was an inversion of two genes detected within the region that codes for five mitochondrial tRNAs (tryptophan, alanine, asparagine, cysteine, and tyrosine), known as the WANCY region, resulting in the order of these genes to appear as WACNY (Figure 5a). Their order was determined by Mitfi annotation of the 176 HiFi reads. All of the reads had enough tRNAs to affirm the WACNY order; 174 encompassed all of these 5 tRNA genes, and the other two reads began with CNY and NY, also indicating the WACNY gene order. There were also 135 HiFi read excerpts that encompassed the *Pro* tRNA gene, the entire control region (CR), and the *Phe* tRNA gene from which we determined the CR lengths (excluding the *Pro* and *Phe* sequences). The length of the CR ranged from 2,620 to 6,544 bp with a mean of 4,243 bp (median = 4,317 bp) (Figure 5b). Of the 135 sequences, 130 had a 60 bp repeat beginning after the *Pro* tRNA (consensus sequence: CCCCCCGTTCGGGCTTTGCTTAAGTCCATGCTAATATATTTCCTTTTTTTTTCGTCCGCA), and the other 5 reads had similar repeats. This sequence, or a 1 to 4 nucleotide indel or SNP variation of it, was repeated just under twice up to 69 times in each read. A goose hairpin sequence (Quinn & Wilson 1993), in this case C7TAC7, was found in 133 of the 135 CR sequences (the two others had C_7_TCAC_7_ and C_7_TAC_4_CAC_8_). All of the hairpins started between 350-360 bp from the end of the CR region (the base before the start of tRNA *Phe*), with 105 of them 353 bp or 354 bp from the end (Figure 5a).

**Figure 5.**
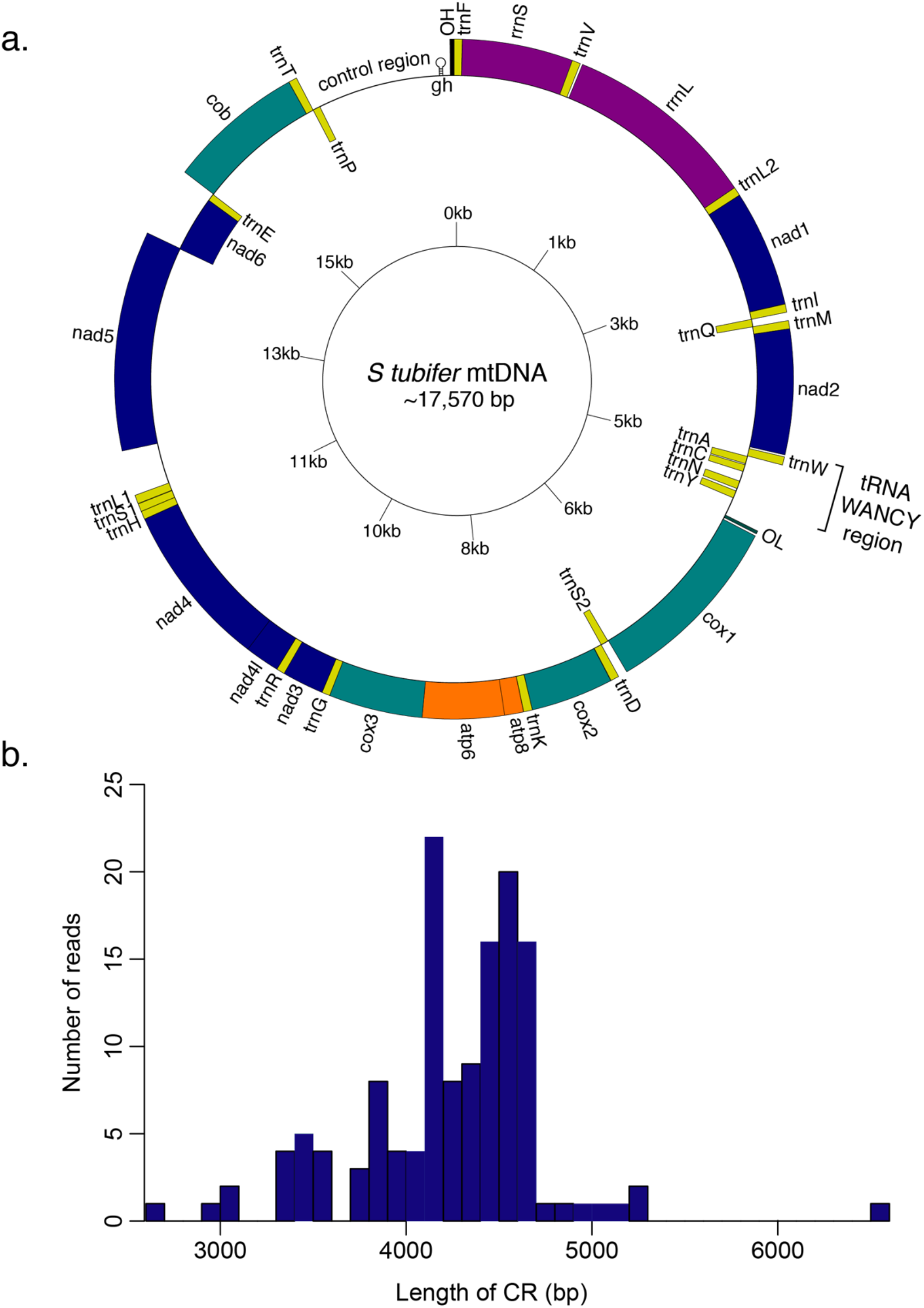
**a)** Gene map of the complete mitogenome of *Siphamia tubifer*. All genes are labelled in addition to the tRNA WANCY region, the control region, and the approximate location of the goose hairpin (gh) within the control region. **b)** Histogram depicting heteroplasmy in the length of the control region observed for the HiFi sequence reads spanning the entire region.

The maximum likelihood phylogeny based on whole mitochondrial sequences (excluding the control region) indicates that *Siphamia tubifer* is divergent from rest of the Apogonidae but a member of the Apogonoidei clade, which also contains the *Kurtus* genus and is sister to the Gobioidei clade (Ghezelayagh *et al.* 2021) (Figure 6). The placement of *S. tubifer* as divergent from the other apogonids is also observed when analyzing a concatenation of several mitochondrial genes, excluding the WANCY tRNA genes (Figure S1). An analysis of *COI* on its own, however, does not align with the other tree topologies, nesting *Siphamia tubifer* within the cardinalfishes, sister to *Ostorhinchus novemfasciatus,* although with low bootstrap support (Figure S1).

**Figure 6.**
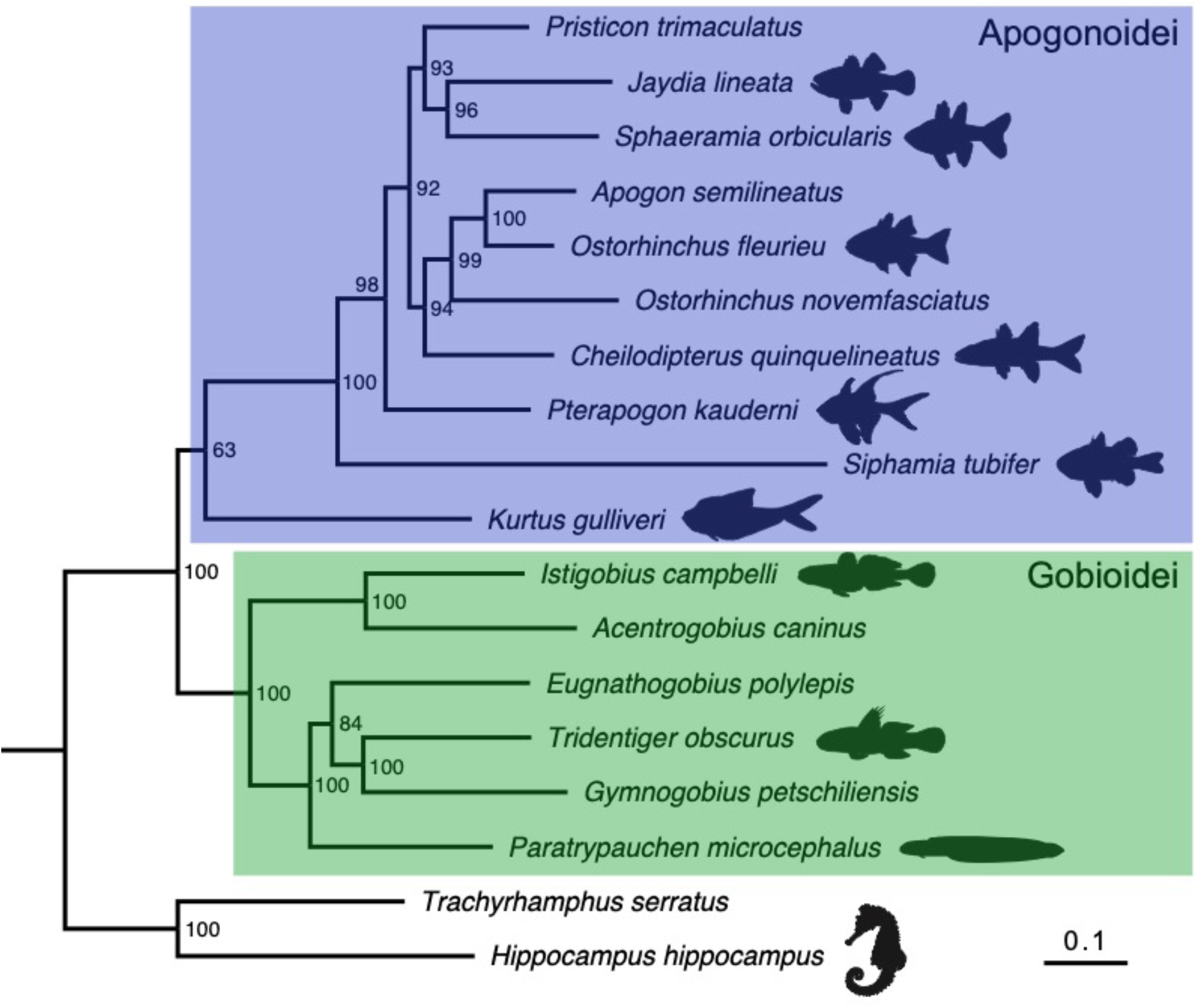
Maximum likelihood tree depicting the phylogenetic relationships of the cardinalfish species for which there is whole mitochondrial genome data available, including *Siphamia tubifer* from this study, in relation to another member of the Apogonoidei clade and several species of gobies. Two Syngnathiformes species are included as an outgroup. The relationships are based on whole mitochondrial DNA sequences excluding the control region using the GTR+F+I+G4 model of substitution. Bootstrap support values (500 replicates) are listed at the nodes. The scale bar indicates nucleotide substitutions per site. The GenBank accession numbers for each species is listed in Table S3.

## Discussion

Combining PacBio HiFi sequencing with Hi-C technology, we assembled a high-quality, chromosome-level genome for the symbiotically luminous cardinalfish *Siphamia tubifer*. The BUSCO score of 99% completeness indicates that this is a near complete genome and will thus serve as a valuable resource for future studies, particularly as the bioluminescent symbiosis between *S. tubifer* and *P. mandapamensis* continues to develop as a tractable, binary model system for symbiosis research. This is only the second cardinalfish genome assembly to date, and our comparison of the two indicates there is significant synteny between them, despite the divergence of *S. tubifer* from the rest of the family. An additional comparison to a more distant genome belonging to the sister order Gobiiformes, revealed a merging of two chromosomes resulting in one fewer chromosome in the cardinalfish genomes. This chromosome fusion also supports the lack of a chromosome 23 in the labelling of the *S. orbicularis* chromosomes, which were named based on synteny with the medaka genome. Determining whether this is a common feature of all cardinalfish genomes and when this merge occurred would require additional chromosomal-level genome assemblies for species in the two orders.

The orthologous cluster analysis between *S. tubifer,* a non-luminous cardinalfish species, and a more distant relative in the Gobiiformes order revealed 710 protein clusters unique to *S. tubifer*. Among these unique clusters, there could be candidate genes that play an important role in the bioluminescent symbiosis. In particular, several clusters were assigned GO terms with functions relating to the immune system and interactions between organisms. There were also several GO terms relating to virus-host interactions that were significantly enriched in the unique protein clusters identified for *S. tubifer*. Although it would require further investigation, the genes involved could play a role in the fish’s interaction with the luminous symbiont. Also of note, visceral muscle development was enriched in the set of unique *S. tubifer* genes. The disc-shaped light organ of *S. tubifer* develops as an outcropping of the gut epithelia and is covered by a lens composed of bundles of transparent muscle tissue on its ventral side. Translucent musculature known as the diffuser also runs along the ventral surface of the fish from the caudal peduncle to the throat, which acts to disperse the light produced by the bacteria inside the light organ (Iwai 1971, Dunlap & Nakamura 2011). The genes associated with the visceral muscle development clusters enriched in *S. tubifer* could be responsible for the development of the light organ and its associated musculature. Additionally, the second largest protein cluster unique to *S. tubifer* was associated with visual perception. There is large overlap in the genes expressed in the light organ and eyes of the symbiotically luminous squid, *Euprymna scolopes,* and the genetic signature specific to the squid light organ includes crystallin and reflectin genes, both of which are typical features of the vertebrate eye (Belcaid *et al.* 2019). Thus, the large cluster of proteins relating to visual perception unique to *S. tubifer* could similarly be related to genes associated with the light organ, such as crystallin and reflection genes. Future studies are needed to determine if the same overlap between the light organ and eye transcripts exist for *S. tubifer.*

A byproduct of HiFi reads for vertebrates, and many bilaterians, is the large percentage of mitogenome sequence captured in an individual read (Formenti *et al.* 2021). These genomes are typically in the range ~16,000 to ~22,000 bp, and their GENBANK annotations canonically start at tRNA *Phe* and end at the control region, which makes them amenable to discover reordering, duplicated regions leading to pseudo-genes, duplicated control regions, control region repeats, and heteroplasmy associated with those and other elements of the mitogenome. With 176 mitochondrial HiFi reads in this study, each a significant percentage of the mitogenome, we were able to determine the unique WACNY ordering of the vertebrate WANCY region of tRNAs of this individual not reported in 3,034 MitoFish website annotations (downloaded June 3, 2021 from http://mitofish.aori.u-tokyo.ac.jp). However, mitochondrial gene-order rearrangements have been observed multiple times in teleost fishes (e.g. Inoue et al 2003, Poulsen et al. 2013), including rearrangements in the WANCY region. For example, a WNCAY tRNA gene order was observed for the blunt snout smooth-head *Xenodermichthys copei* and was most parsimoniously explained by duplications of parts of the mt genome with subsequent deletions (Poulsen et al. 2019). Additional sequencing of the mitochondrial genomes of more *S. tubifer* specimens as well as other *Siphamia* species would indicate whether the WACNY gene order observed in this study is unique to this individual or is a common feature of this species or genus.

PacBio HiFi reads have lower error rates than earlier long read technology, though of course errors exist and homopolymer miscalls are a known class of these. It is likely that some of the differences in the 135 reads that incorporated an entire control region, flanked by the expected tRNAs, come from sequencing error and not the control region itself. However, given that the final part of the CR, which is not repetitive, varies much less in length and sequence than the repeat section of the CR, and the fact that there are almost 4,000 bp between the smallest and largest representations (and over 2200 bp between second smallest and second largest), repeat expansion and/or contraction is likely occurring in the mitochondria of this organism. Heteroplasmy in the length of the control region has been documented for other fish species, including the three-spined stickleback (Stärner *et al.* 2004), two species of sardines (Samonte *et al.* 2000), the flatfish *Platichthys flesus* (Hoarau *et al.* 2002), and several sturgeon species (Ludwig *et al.* 2000). Such variability in the copy number of tandem repeats in the control region could be a more common occurrence that has been overlooked with previous sequencing approaches. Thus, the ability of HiFi reads to reveal heteroplasmy in the mitogenome could lead to increased observations of this phenomenon in other organisms (e.g. Formenti *et al.* 2021) as HiFi sequencing becomes more widely implemented. Importantly, variability in the length of the control region has previously been used as a genetic marker to discriminate between species (e.g. Faber & Stepien 1998, Turanov *et al.* 2019). If heteroplasmy in the control region is a more common occurrence, then its use as a marker could be erroneous in many cases.

The phylogenetic analysis based on whole mitochondrial genome sequences indicated that *S. tubifer* is divergent from the other members of the cardinalfish family Apogonidae, a placement previously supported and estimated to have occurred approximately 50 million years ago (Thacker 2014). The evolutionary relationship of *S. tubifer* as sister to the rest of the cardinalfishes raises the possibility that the bioluminescent symbiosis with *P. mandapamensis* played a role in the host’s initial divergence and speciation from a common ancestor. The acquisition of bacterial endosymbionts was proposed nearly a century ago as a primary mechanism by which new species can arise (Wallin 1927), and speciation by symbiosis has since been documented, primarily for several insect hosts (Brucker & Bordenstein 2012). Future studies identifying host genes involved in the *S. tubifer*-*P. mandapamensis* symbiosis will help determine whether there is any evidence that the symbiosis played a role in speciation for *Siphamia*. The high-quality genome assembly for *S. tubifer* presented here will serve as a valuable resource for both the study of the evolution of symbiotic bioluminescence in fishes as well as the functional genomics of the symbiosis, further establishing the *S. tubifer*-*P. mandapamensis* association as a tractable model for the study of vertebrate-bacteria interactions and microbial symbiosis more broadly.

## Data Accessibility

Genome assembly and associated sequencing data are available under NCBI Bioproject PRJNA736963.

## Author Contributions

ALG conceived of the project and secured funding for the work. ALG carried out tissue dissections and AWL preformed the high molecular weight DNA extractions and HI-C library preparations. JBH carried out the genome assembly and associated bioinformatics. ALG performed the phylogenetic analyses and data analyses. ALG and JBH contributed to the discussion and interpretation of the results and writing of the manuscript. All authors approve of the submitted version of this manuscript.

