## Supplemental Materials for "Chromosomal-level genome assembly of the bioluminescent cardinalfish *Siphamia tubifer*, an emerging model for symbiosis research"

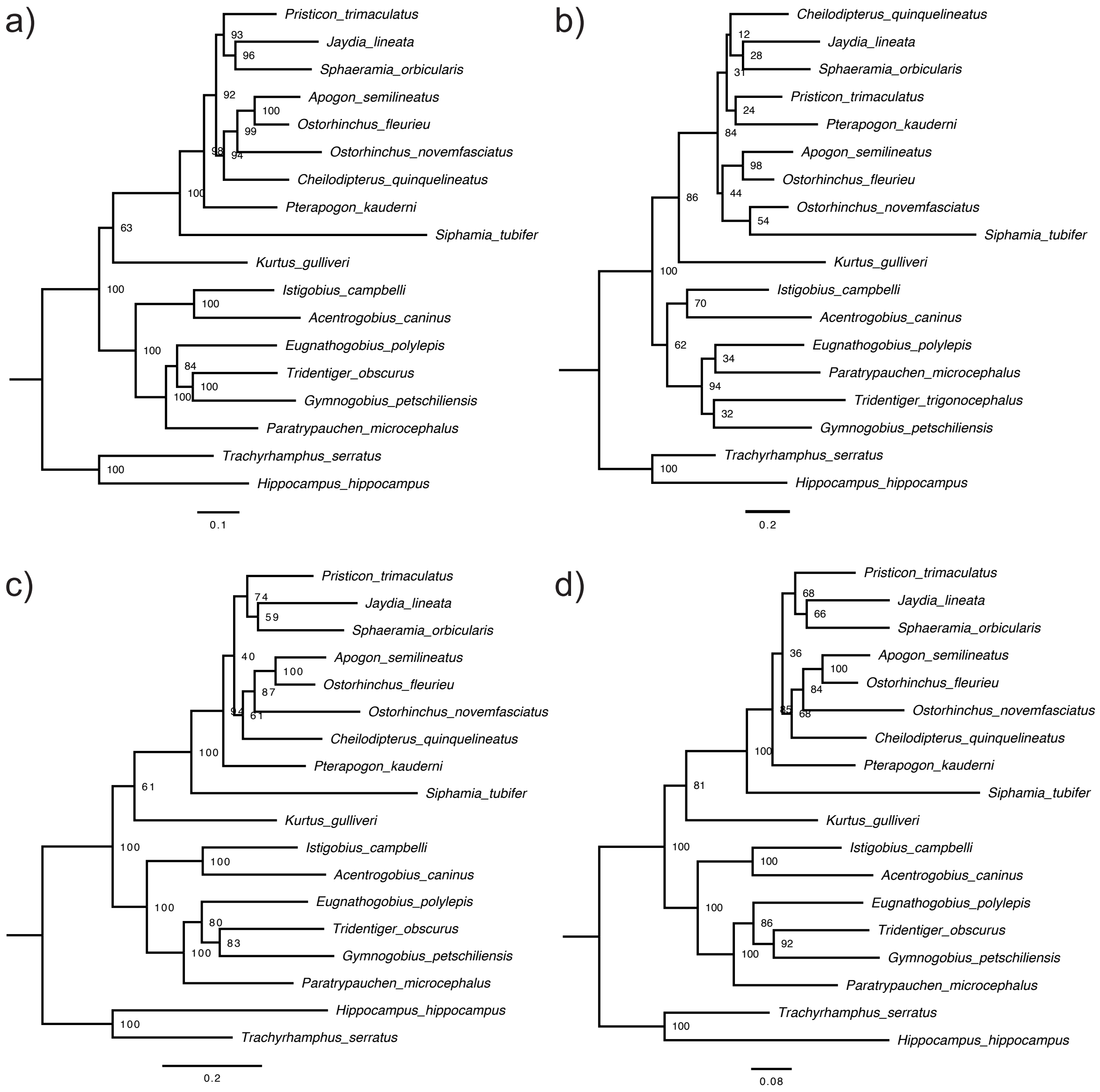


**Figure S1**. Maximum likelihood trees depicting the phylogenetic relationships between several Gobiiformes and Kurtiformes species for which whole mitochondrial sequences are available, including *S. tubifer* from this study. Models based on sequences from: a) whole mitochondrial genomes excluding the control region b) *COI* only c) concatenated mitochondrial genes: *COI*, *ND4*, 16S, and *cytB*, and d) the same genes as c) excluding *cytB*. The substitution model used was GTR+F+I+G4. Bootstrap support values are included at the nodes. Scale bars indicate nucleotide substitutions per site.

**Table S1**. List of 506 of the 701 total protein clusters unique to *Siphamia tubifer* assigned GO annotations**.** The number of proteins in each cluster is indicated as is the associated SWISS-PROT ID.

| Cluster ID | Proteins | SWISS-PROT ID | GO annotation |
|---|---|---|---|
| cluster20 | 70 | P20825 | GO:0015074; P:DNA integration; IEA:InterPro |
| cluster50 | 41 | Q52M02 | GO:0007601; P:visual perception; TAS:ProtInc |
| cluster71 | 32 | P0CT41 | GO:0006310; P:DNA recombination; IEA:UniProtKB-KW |
| cluster119 | 25 | Q80Z10 | GO:0015031; P:protein transport; IEA:UniProtKB-KW |
| cluster146 | 23 | P27401 | GO:0075732; P:viral penetration into host nucleus; IEA:UniProtKB-KW |
| cluster162 | 22 | Q03278 | GO:0003964; F:RNA-directed DNA polymerase activity; IEA:UniProtKB-KW |
| cluster163 | 22 | O95125 | GO:0006366; P:transcription by RNA polymerase II; TAS:ProtInc |
| cluster181 | 21 | A1X283 | GO:0006801; P:superoxide metabolic process; IDA:UniProtKB |
| cluster224 | 19 | Q9M2N5 | GO:0009791; P:post-embryonic development; IMP:TAIR |
| cluster275 | 17 | O92815 | GO:0006310; P:DNA recombination; IEA:UniProtKB-KW |
| cluster507 | 12 | O92815 | GO:0006310; P:DNA recombination; IEA:UniProtKB-KW |
| cluster516 | 12 | Q68EI0 | GO:0006364; P:rRNA processing; IBA:GO_Central |
| cluster717 | 10 | A0MSJ1 | GO:0001501; P:skeletal system development; IGI:ZFIN |
| cluster737 | 10 | A1X283 | GO:0006801; P:superoxide metabolic process; IDA:UniProtKB |
| cluster868 | 9 | O60290 | GO:0006355; P:regulation of transcription, DNA-templated; IEA:InterPro |
| cluster1140 | 8 | Q95218 | GO:0015031; P:protein transport; IEA:UniProtKB-KW |
| cluster1146 | 8 | Q14258 | GO:0016032; P:viral process; IEA:UniProtKB-KW |
| cluster1147 | 8 | P43353 | GO:0030148; P:sphingolipid biosynthetic process; TAS:Reactome |
| cluster1127 | 8 | E1C5V0 | GO:0043516; P:regulation of DNA damage response, signal transduction by p53 class mediator; ISS:UniProtKB |
| cluster1165 | 8 | O96006 | GO:0045944; P:positive regulation of transcription by RNA polymerase II; IDA:NTNU_SB |
| cluster1596 | 7 | Q1RLU1 | GO:0010212; P:response to ionizing radiation; ISS:UniProtKB |
| cluster1552 | 7 | Q9HCI6 | GO:0016925; P:protein sumoylation; IDA:UniProtKB |
| cluster2199 | 6 | Q7TN75 | GO:0001890; P:placenta development; IMP:MGI |
| cluster2756 | 6 | P02264 | GO:0051673; P:membrane disruption in other organism; IDA:AgBase |
| cluster3243 | 5 | P85521 | GO:0006953; P:acute-phase response; IEA:UniProtKB-KW |
| cluster3432 | 5 | Q9UPY6 | GO:0008360; P:regulation of cell shape; IMP:UniProtKB |
| cluster3290 | 5 | P51574 | GO:0015031; P:protein transport; IEA:UniProtKB-KW |
| cluster3212 | 5 | P10394 | GO:0015074; P:DNA integration; IEA:InterPro |

| | | | |
|---|---|---|---|
| cluster3282 | 5 | P03360 | GO:0044826; P:viral genome integration into host DNA; IEA:UniProtKB-KW |
| cluster3245 | 5 | O75132 | GO:0046983; F:protein dimerization activity; IEA:InterPro |
| cluster3295 | 5 | F8RKW0 | GO:0090729; F:toxin activity; IEA:UniProtKB-KW |
| cluster7229 | 4 | Q9QXS4 | GO:0001702; P:gastrulation with mouth forming second; NAS:UniProtKB |
| cluster5270 | 4 | Q3B8D4 | GO:0003677; F:DNA binding; IEA:UniProtKB-KW |
| cluster5259 | 4 | P29401 | GO:0005999; P:xylulose biosynthetic process; TAS:Reactome |
| cluster5567 | 4 | P11369 | GO:0006310; P:DNA recombination; IEA:UniProtKB-KW |
| cluster5367 | 4 | P61648 | GO:0006493; P:protein O-linked glycosylation; IEA:Ensembl |
| cluster5315 | 4 | Q9NU92 | GO:0007165; P:signal transduction; TAS:ProtInc |
| cluster7149 | 4 | Q5F4A1 | GO:0007275; P:multicellular organism development; IEA:UniProtKB-KW |
| cluster5442 | 4 | Q99P25 | GO:0007283; P:spermatogenesis; IEA:UniProtKB-KW |
| cluster5543 | 4 | Q1RLU1 | GO:0010212; P:response to ionizing radiation; ISS:UniProtKB |
| cluster7123 | 4 | P04323 | GO:0015074; P:DNA integration; IEA:InterPro |
| cluster5430 | 4 | Q14534 | GO:0016126; P:sterol biosynthetic process; IBA:GO_Central |
| cluster5477 | 4 | B1WC10 | GO:0032185; P:septin cytoskeleton organization; ISS:UniProtKB |
| cluster5275 | 4 | P11260 | GO:0032197; P:transposition, RNA-mediated; IMP:UniProtKB |
| cluster5482 | 4 | Q9QXR7 | GO:0042060; P:wound healing; IBA:GO_Central |
| cluster5471 | 4 | Q7ZXV5 | GO:0043516; P:regulation of DNA damage response, signal transduction by p53 class mediator; ISS:UniProtKB |
| cluster7231 | 4 | Q96TG0 | GO:0045995; P:regulation of embryonic development; IEA:InterPro |
| cluster7076 | 4 | Q9HAC7 | GO:0047369; F:succinate-hydroxymethylglutarate CoA-transferase activity; IDA:UniProtKB |
| cluster5219 | 4 | Q19546 | GO:0061820; P:telomeric D-loop disassembly; IBA:GO_Central |
| cluster7072 | 4 | Q09811 | GO:0070914; P:UV-damage excision repair; IMP:PomBase |
| cluster5137 | 4 | O94885 | GO:1900044; P:regulation of protein K63-linked ubiquitination; IDA:MGI |
| cluster9379 | 3 | Q91WP0 | GO:0001867; P:complement activation, lectin pathway; IDA:MGI |
| cluster9382 | 3 | Q4U4S6 | GO:0003281; P:ventricular septum development; IMP:MGI |
| cluster9423 | 3 | Q04202 | GO:0006313; P:transposition, DNA-mediated; IEA:InterPro |
| cluster9451 | 3 | Q9NBX4 | GO:0006313; P:transposition, DNA-mediated; IMP:UniProtKB |
| cluster12652 | 3 | Q61510 | GO:0006511; P:ubiquitin-dependent protein catabolic process; ISO:MGI |
| cluster9464 | 3 | Q9UN72 | GO:0007399; P:nervous system development; TAS:ProtInc |
| cluster12656 | 3 | P55067 | GO:0007417; P:central nervous system development; IBA:GO_Central |
| cluster9405 | 3 | Q5KQS3 | GO:0008191; F:metalloendopeptidase inhibitor activity; IEA:UniProtKB-KW |

| | | | |
|---|---|---|---|
| cluster12672 | 3 | Q8BV66 | GO:0009617; P:response to bacterium; IEP:MGI |
| cluster9435 | 3 | Q1RLU1 | GO:0010212; P:response to ionizing radiation; ISS:UniProtKB |
| cluster12653 | 3 | P36514 | GO:0015020; F:glucuronosyltransferase activity; IEA:UniProtKB-EC |
| cluster9449 | 3 | Q9UKJ4 | GO:0016032; P:viral process; IEA:UniProtKB-KW |
| cluster9469 | 3 | Q7LHG5 | GO:0032197; P:transposition, RNA-mediated; ISS:SGD |
| cluster12646 | 3 | Q9PTM4 | GO:0034765; P:regulation of ion transmembrane transport; IEA:UniProtKB-KW |
| cluster9447 | 3 | Q92820 | GO:0046900; P:tetrahydrofolylpolyglutamate metabolic process; IBA:GO_Central |
| cluster9444 | 3 | Q96DH6 | GO:0048864; P:stem cell development; IEA:Ensembl |
| cluster9365 | 3 | Q9Y5F6 | GO:0050808; P:synapse organization; IEA:Ensembl |
| cluster9402 | 3 | P02264 | GO:0051673; P:membrane disruption in other organism; IDA:AgBase |
| cluster9367 | 3 | P20693 | GO:0051770; P:positive regulation of nitric-oxide synthase biosynthetic process; ISO:MGI |
| cluster9431 | 3 | F1QH17 | GO:0055114; P:oxidation-reduction process; ISS:UniProtKB |
| cluster9430 | 3 | E9Q401 | GO:0086005; P:ventricular cardiac muscle cell action potential; IMP:BHF-UCL |
| cluster12657 | 3 | O54898 | GO:0086018; P:SA node cell to atrial cardiac muscle cell signaling; ISS:BHF-UCL |
| cluster9436 | 3 | Q8R4F1 | GO:0099560; P:synaptic membrane adhesion; IDA:SynGO |
| cluster9369 | 3 | Q16820 | GO:1901998; P:toxin transport; IEA:Ensembl |
| cluster18310 | 2 | Q66I24 | GO:0000050; P:urea cycle; ISS:UniProtKB |
| cluster19257 | 2 | Q9H6P0 | GO:0000122; P:negative regulation of transcription by RNA polymerase II; IDA:BHF-UCL |
| cluster18344 | 2 | Q86VZ6 | GO:0000122; P:negative regulation of transcription by RNA polymerase II; IDA:MGI |
| cluster19277 | 2 | Q86VZ6 | GO:0000122; P:negative regulation of transcription by RNA polymerase II; IDA:MGI |
| cluster18351 | 2 | Q6P0B1 | GO:0000381; P:regulation of alternative mRNA splicing, via spliceosome; IBA:GO_Central |
| cluster18358 | 2 | Q7T2T1 | GO:0000381; P:regulation of alternative mRNA splicing, via spliceosome; IBA:GO_Central |
| cluster19259 | 2 | Q7T2T1 | GO:0000381; P:regulation of alternative mRNA splicing, via spliceosome; IBA:GO_Central |
| cluster19182 | 2 | A1A4K8 | GO:0000398; P:mRNA splicing, via spliceosome; IBA:GO_Central |
| cluster18403 | 2 | Q8QGX4 | GO:0000723; P:telomere maintenance; IBA:GO_Central |
| cluster19154 | 2 | Q9Y6X0 | GO:0000981; F:DNA-binding transcription factor activity, RNA polymerase II-specific; ISM:NTNU_SB |
| cluster18275 | 2 | Q8VHG2 | GO:0001570; P:vasculogenesis; IMP:MGI |

| | | | |
|---|---|---|---|
| cluster18319 | 2 | E9PCK8 | GO:0001657; P:ureteric bud development; ISS:BHF-UCL |
| cluster18409 | 2 | P48645 | GO:0001659; P:temperature homeostasis; IEA:Ensembl |
| cluster18490 | 2 | Q9JI18 | GO:0001701; P:in utero embryonic development; IMP:MGI |
| cluster18317 | 2 | Q96II1 | GO:0001764; P:neuron migration; IBA:GO_Central |
| cluster18350 | 2 | Q96II1 | GO:0001764; P:neuron migration; IBA:GO_Central |
| cluster19162 | 2 | Q61137 | GO:0001764; P:neuron migration; IDA:MGI |
| cluster18460 | 2 | P98064 | GO:0001867; P:complement activation, lectin pathway; IMP:UniProtKB |
| cluster19296 | 2 | Q9Z2W9 | GO:0001919; P:regulation of receptor recycling; ISO:MGI |
| cluster19326 | 2 | O15399 | GO:0001964; P:startle response; IEA:Ensembl |
| cluster18303 | 2 | Q96SG3 | GO:0001967; P:suckling behavior; IEA:Ensembl |
| cluster19243 | 2 | Q96SG3 | GO:0001967; P:suckling behavior; IEA:Ensembl |
| cluster19400 | 2 | A1XQX1 | GO:0002040; P:sprouting angiogenesis; IMP:ZFIN |
| cluster19165 | 2 | Q56JV1 | GO:0002181; P:cytoplasmic translation; ISS:UniProtKB |
| cluster18456 | 2 | A6QLK5 | GO:0002437; P:inflammatory response to antigenic stimulus; ISS:UniProtKB |
| cluster19261 | 2 | P13789 | GO:0003009; P:skeletal muscle contraction; IBA:GO_Central |
| cluster18346 | 2 | P22004 | GO:0003323; P:type B pancreatic cell development; IDA:BHF-UCL |
| cluster19205 | 2 | Q04906 | GO:0003323; P:type B pancreatic cell development; IEA:Ensembl |
| cluster18324 | 2 | P32018 | GO:0003429; P:growth plate cartilage chondrocyte morphogenesis; IBA:GO_Central |
| cluster19244 | 2 | P13944 | GO:0003429; P:growth plate cartilage chondrocyte morphogenesis; IBA:GO_Central |
| cluster19216 | 2 | A6H7H1 | GO:0003723; F:RNA binding; IEA:UniProtKB-KW |
| cluster18383 | 2 | Q8MJU9 | GO:0003774; F:motor activity; IEA:InterPro |
| cluster18477 | 2 | Q00962 | GO:0003964; F:RNA-directed DNA polymerase activity; IEA:UniProtKB-KW |
| cluster18401 | 2 | Q7SIG3 | GO:0004252; F:serine-type endopeptidase activity; IEA:InterPro |
| cluster18282 | 2 | B6ZK76 | GO:0004908; F:interleukin-1 receptor activity; IEA:InterPro |
| cluster18448 | 2 | Q8NGI2 | GO:0004984; F:olfactory receptor activity; IBA:GO_Central |
| cluster18504 | 2 | P51856 | GO:0005198; F:structural molecule activity; IEA:InterPro |
| cluster18365 | 2 | P98157 | GO:0005509; F:calcium ion binding; IEA:InterPro |
| cluster18425 | 2 | O02751 | GO:0005634; C:nucleus; IEA:UniProtKB-SubCell |
| cluster18533 | 2 | Q6DDT5 | GO:0005737; C:cytoplasm; ISS:UniProtKB |
| cluster19208 | 2 | A0JM08 | GO:0005874; C:microtubule; IEA:UniProtKB-KW |
| cluster18498 | 2 | Q2M3C6 | GO:0005886; C:plasma membrane; IDA:HPA |
| cluster19283 | 2 | F1P4W9 | GO:0005886; C:plasma membrane; IEA:UniProtKB-KW |
| cluster19276 | 2 | Q6IPM2 | GO:0005929; C:cilium; TAS:Reactome |

| | | | |
|---|---|---|---|
| cluster18421 | 2 | B5DGQ7 | GO:0006096; P:glycolytic process; IEA:UniProtKB-UniPathway |
| cluster18304 | 2 | Q7TQ07 | GO:0006261; P:DNA-dependent DNA replication; ISO:MGI |
| cluster19339 | 2 | Q7TQ07 | GO:0006261; P:DNA-dependent DNA replication; ISO:MGI |
| cluster19192 | 2 | O93309 | GO:0006275; P:regulation of DNA replication; ISS:UniProtKB |
| cluster18356 | 2 | Q92089 | GO:0006304; P:DNA modification; IEA:UniProtKB-KW |
| cluster18523 | 2 | P0CT41 | GO:0006310; P:DNA recombination; IEA:UniProtKB-KW |
| cluster18377 | 2 | Q9NBX4 | GO:0006313; P:transposition, DNA-mediated; IMP:UniProtKB |
| cluster19354 | 2 | Q9NBX4 | GO:0006313; P:transposition, DNA-mediated; IMP:UniProtKB |
| cluster19179 | 2 | Q9NTU8 | GO:0006351; P:transcription, DNA-templated; IEA:InterPro |
| cluster18245 | 2 | P40645 | GO:0006355; P:regulation of transcription, DNA-templated; IDA:MGI |
| cluster18441 | 2 | Q9GL32 | GO:0006357; P:regulation of transcription by RNA polymerase II; IBA:GO_Central |
| cluster19297 | 2 | Q86UJ9 | GO:0006366; P:transcription by RNA polymerase II; TAS:ProtInc |
| cluster18357 | 2 | P42568 | GO:0006368; P:transcription elongation from RNA polymerase II promoter; TAS:Reactome |
| cluster19327 | 2 | Q9C0J8 | GO:0006369; P:termination of RNA polymerase II transcription; TAS:Reactome |
| cluster18261 | 2 | Q92005 | GO:0006414; P:translational elongation; IBA:GO_Central |
| cluster19349 | 2 | Q8BWD8 | GO:0006468; P:protein phosphorylation; IBA:GO_Central |
| cluster18309 | 2 | Q15208 | GO:0006468; P:protein phosphorylation; IDA:UniProtKB |
| cluster18408 | 2 | Q8WX83 | GO:0006468; P:protein phosphorylation; IMP:UniProtKB |
| cluster18510 | 2 | Q8WX83 | GO:0006468; P:protein phosphorylation; IMP:UniProtKB |
| cluster19310 | 2 | O62830 | GO:0006470; P:protein dephosphorylation; ISS:UniProtKB |
| cluster19196 | 2 | E6ZGB4 | GO:0006482; P:protein demethylation; ISS:UniProtKB |
| cluster19273 | 2 | E6ZGB4 | GO:0006482; P:protein demethylation; ISS:UniProtKB |
| cluster18535 | 2 | Q6P9A2 | GO:0006493; P:protein O-linked glycosylation; IDA:UniProtKB |
| cluster19378 | 2 | O08688 | GO:0006508; P:proteolysis; IBA:GO_Central |
| cluster19313 | 2 | Q9ULZ9 | GO:0006508; P:proteolysis; TAS:ParkinsonsUK-UCL |
| cluster18414 | 2 | O70263 | GO:0006511; P:ubiquitin-dependent protein catabolic process; IDA:MGI |
| cluster19207 | 2 | Q14974 | GO:0006610; P:ribosomal protein import into nucleus; IDA:UniProtKB |
| cluster19223 | 2 | F1QXD3 | GO:0006612; P:protein targeting to membrane; IBA:GO_Central |
| cluster18355 | 2 | P50416 | GO:0006641; P:triglyceride metabolic process; IEA:Ensembl |
| cluster18345 | 2 | Q9HBU6 | GO:0006646; P:phosphatidylethanolamine biosynthetic process; IDA:UniProtKB |
| cluster19357 | 2 | Q9D4V0 | GO:0006646; P:phosphatidylethanolamine biosynthetic process; ISO:MGI |
| cluster19345 | 2 | Q9JJY3 | GO:0006684; P:sphingomyelin metabolic process; IDA:MGI |

| | | | |
|---|---|---|---|
| cluster19409 | 2 | P79386 | GO:0006694; P:steroid biosynthetic process; ISS:HGNC |
| cluster19294 | 2 | Q86YH6 | GO:0006744; P:ubiquinone biosynthetic process; IDA:HGNC |
| cluster19312 | 2 | P30561 | GO:0006805; P:xenobiotic metabolic process; TAS:MGI |
| cluster19334 | 2 | Q3YAW7 | GO:0006811; P:ion transport; IEA:UniProtKB-KW |
| cluster18393 | 2 | Q99250 | GO:0006814; P:sodium ion transport; TAS:ProtInc |
| cluster18532 | 2 | Q99250 | GO:0006814; P:sodium ion transport; TAS:ProtInc |
| cluster18388 | 2 | P53992 | GO:0006886; P:intracellular protein transport; IEA:InterPro |
| cluster19263 | 2 | Q9NZR2 | GO:0006898; P:receptor-mediated endocytosis; TAS:ProtInc |
| cluster18306 | 2 | Q8WU76 | GO:0006904; P:vesicle docking involved in exocytosis; IEA:InterPro |
| cluster18497 | 2 | A0JMW6 | GO:0006915; P:apoptotic process; IEA:UniProtKB-KW |
| cluster19169 | 2 | O09127 | GO:0006929; P:substrate-dependent cell migration; IDA:UniProtKB |
| cluster19362 | 2 | Q9UIT1 | GO:0006939; P:smooth muscle contraction; TAS:ProtInc |
| cluster18470 | 2 | Q2VLH6 | GO:0006953; P:acute-phase response; IEA:UniProtKB-KW |
| cluster19226 | 2 | Q2VLG6 | GO:0006953; P:acute-phase response; IEA:UniProtKB-KW |
| cluster18431 | 2 | P98093 | GO:0006954; P:inflammatory response; IEA:UniProtKB-KW |
| cluster19346 | 2 | Q9WU60 | GO:0006979; P:response to oxidative stress; ISO:MGI |
| cluster18392 | 2 | Q5KR47 | GO:0007015; P:actin filament organization; IBA:GO_Central |
| cluster18273 | 2 | Q7Z668 | GO:0007018; P:microtubule-based movement; IBA:GO_Central |
| cluster19161 | 2 | E9PYY5 | GO:0007018; P:microtubule-based movement; IBA:GO_Central |
| cluster19194 | 2 | Q7Z668 | GO:0007018; P:microtubule-based movement; IBA:GO_Central |
| cluster19228 | 2 | Q9NQ86 | GO:0007051; P:spindle organization; IMP:UniProtKB |
| cluster18243 | 2 | Q9DDD0 | GO:0007155; P:cell adhesion; IEA:UniProtKB-KW |
| cluster18339 | 2 | A1XQY1 | GO:0007155; P:cell adhesion; IEA:UniProtKB-KW |
| cluster18361 | 2 | A2AVA0 | GO:0007155; P:cell adhesion; IEA:UniProtKB-KW |
| cluster18482 | 2 | P0C6B8 | GO:0007155; P:cell adhesion; IEA:UniProtKB-KW |
| cluster18318 | 2 | P24503 | GO:0007156; P:homophilic cell adhesion via plasma membrane adhesion molecules; IBA:GO_Central |
| cluster18458 | 2 | P24503 | GO:0007156; P:homophilic cell adhesion via plasma membrane adhesion molecules; IBA:GO_Central |
| cluster19398 | 2 | Q5DRB1 | GO:0007156; P:homophilic cell adhesion via plasma membrane adhesion molecules; IEA:InterPro |
| cluster18487 | 2 | P28472 | GO:0007165; P:signal transduction; IBA:GO_Central |
| cluster19338 | 2 | Q13191 | GO:0007165; P:signal transduction; IBA:GO_Central |

| | | | |
|---|---|---|---|
| cluster18395 | 2 | Q8TCX5 | GO:0007165; P:signal transduction; IEA:InterPro |
| cluster18491 | 2 | A6NI28 | GO:0007165; P:signal transduction; IEA:InterPro |
| cluster19174 | 2 | Q6DRG7 | GO:0007165; P:signal transduction; IEA:InterPro |
| cluster19240 | 2 | Q6TLK4 | GO:0007165; P:signal transduction; IEA:InterPro |
| cluster19350 | 2 | Q61302 | GO:0007165; P:signal transduction; IEA:InterPro |
| cluster18269 | 2 | Q9UJF2 | GO:0007165; P:signal transduction; TAS:ProtInc |
| cluster18437 | 2 | Q92915 | GO:0007165; P:signal transduction; TAS:ProtInc |
| cluster18544 | 2 | Q9NU92 | GO:0007165; P:signal transduction; TAS:ProtInc |
| cluster19199 | 2 | Q8WUQ3 | GO:0007165; P:signal transduction; TAS:ProtInc |
| cluster19211 | 2 | O14514 | GO:0007165; P:signal transduction; TAS:ProtInc |
| cluster19363 | 2 | O15484 | GO:0007165; P:signal transduction; TAS:ProtInc |
| cluster18509 | 2 | O97814 | GO:0007166; P:cell surface receptor signaling pathway; IEA:InterPro |
| cluster19164 | 2 | Q9UM73 | GO:0007169; P:transmembrane receptor protein tyrosine kinase signaling pathway; IEA:InterPro |
| cluster19181 | 2 | P43026 | GO:0007179; P:transforming growth factor beta receptor signaling pathway; TAS:ProtInc |
| cluster18517 | 2 | P97751 | GO:0007187; P:G protein-coupled receptor signaling pathway, coupled to cyclic nucleotide second messenger; ISO:MGI |
| cluster18283 | 2 | O75899 | GO:0007194; P:negative regulation of adenylate cyclase activity; TAS:ProtInc |
| cluster19250 | 2 | P28335 | GO:0007210; P:serotonin receptor signaling pathway; IMP:UniProtKB |
| cluster19371 | 2 | Q2KI97 | GO:0007218; P:neuropeptide signaling pathway; IBA:GO_Central |
| cluster19332 | 2 | Q8C0Q9 | GO:0007264; P:small GTPase mediated signal transduction; IEA:InterPro |
| cluster19245 | 2 | Q91Z69 | GO:0007266; P:Rho protein signal transduction; IDA:MGI |
| cluster18270 | 2 | Q02084 | GO:0007275; P:multicellular organism development; IEA:UniProtKB-KW |
| cluster18399 | 2 | Q6VNB8 | GO:0007275; P:multicellular organism development; IEA:UniProtKB-KW |
| cluster18436 | 2 | Q90304 | GO:0007275; P:multicellular organism development; IEA:UniProtKB-KW |
| cluster18492 | 2 | Q6DN12 | GO:0007275; P:multicellular organism development; IEA:UniProtKB-KW |
| cluster18541 | 2 | P79777 | GO:0007275; P:multicellular organism development; IEA:UniProtKB-KW |
| cluster19265 | 2 | Q8R508 | GO:0007275; P:multicellular organism development; IEA:UniProtKB-KW |
| cluster19405 | 2 | A0A1L8GSA2 | GO:0007275; P:multicellular organism development; IEA:UniProtKB-KW |
| cluster18417 | 2 | Q6ZWR6 | GO:0007283; P:spermatogenesis; IEA:UniProtKB-KW |
| cluster19176 | 2 | Q08E27 | GO:0007286; P:spermatid development; IEA:Ensembl |
| cluster19254 | 2 | Q9MYW0 | GO:0007338; P:single fertilization; IBA:GO_Central |

| | | | |
|---|---|---|---|
| cluster18472 | 2 | Q5DID0 | GO:0007338; P:single fertilization; IEA:Ensembl |
| cluster18464 | 2 | O57682 | GO:0007367; P:segment polarity determination; IEA:UniProtKB-KW |
| cluster18244 | 2 | O60542 | GO:0007399; P:nervous system development; TAS:ProtInc |
| cluster18364 | 2 | Q9Y5I4 | GO:0007399; P:nervous system development; TAS:ProtInc |
| cluster19255 | 2 | Q63374 | GO:0007416; P:synapse assembly; ISS:BHF-UCL |
| cluster18281 | 2 | Q02566 | GO:0007522; P:visceral muscle development; IMP:MGI |
| cluster19158 | 2 | Q02566 | GO:0007522; P:visceral muscle development; IMP:MGI |
| cluster18241 | 2 | Q8JIR8 | GO:0007601; P:visual perception; IEA:InterPro |
| cluster18522 | 2 | Q8N6F1 | GO:0007601; P:visual perception; IEA:UniProtKB-KW |
| cluster19212 | 2 | Q13402 | GO:0007601; P:visual perception; IMP:UniProtKB |
| cluster18378 | 2 | O73700 | GO:0007605; P:sensory perception of sound; IEA:Ensembl |
| cluster19262 | 2 | Q9UM54 | GO:0007605; P:sensory perception of sound; IEA:UniProtKB-KW |
| cluster19264 | 2 | Q9YH85 | GO:0007605; P:sensory perception of sound; IEA:UniProtKB-KW |
| cluster18302 | 2 | Q01668 | GO:0007605; P:sensory perception of sound; IMP:BHF-UCL |
| cluster19300 | 2 | Q9JK97 | GO:0007605; P:sensory perception of sound; IMP:MGI |
| cluster19287 | 2 | P56696 | GO:0007605; P:sensory perception of sound; TAS:ProtInc |
| cluster19329 | 2 | Q8BHR2 | GO:0007624; P:ultradian rhythm; IEA:InterPro |
| cluster19235 | 2 | Q08C92 | GO:0008033; P:tRNA processing; IEA:UniProtKB-KW |
| cluster19299 | 2 | P81018 | GO:0008061; F:chitin binding; IDA:AgBase |
| cluster19219 | 2 | Q96FA3 | GO:0008063; P:Toll signaling pathway; IBA:GO_Central |
| cluster19210 | 2 | Q96Q29 | GO:0008104; P:protein localization; IBA:GO_Central |
| cluster18341 | 2 | Q8NFP9 | GO:0008104; P:protein localization; IEA:Ensembl |
| cluster18370 | 2 | Q64GL0 | GO:0008154; P:actin polymerization or depolymerization; ISS:UniProtKB |
| cluster18412 | 2 | Q64GL0 | GO:0008154; P:actin polymerization or depolymerization; ISS:UniProtKB |
| cluster18521 | 2 | Q60467 | GO:0008201; F:heparin binding; IEA:UniProtKB-KW |
| cluster19408 | 2 | Q9BY12 | GO:0008270; F:zinc ion binding; IEA:InterPro |
| cluster18473 | 2 | Q5VXN8 | GO:0008360; P:regulation of cell shape; IMP:CAFA |
| cluster19352 | 2 | Q5VXN8 | GO:0008360; P:regulation of cell shape; IMP:CAFA |
| cluster19303 | 2 | Q7YQM1 | GO:0008380; P:RNA splicing; IEA:UniProtKB-KW |
| cluster18423 | 2 | E7FCN8 | GO:0008589; P:regulation of smoothened signaling pathway; ISS:UniProtKB |
| cluster18484 | 2 | Q6NS52 | GO:0009617; P:response to bacterium; IEP:MGI |
| cluster19173 | 2 | Q9UQR0 | GO:0009653; P:anatomical structure morphogenesis; TAS:ProtInc |
| cluster18256 | 2 | P0DM44 | GO:0009755; P:hormone-mediated signaling pathway; IBA:GO_Central |

| | | | |
|---|---|---|---|
| cluster18287 | 2 | Q8WXI1 | GO:0009966; P:regulation of signal transduction; IEA:InterPro |
| cluster18424 | 2 | Q6PJ69 | GO:0010508; P:positive regulation of autophagy; IMP:UniProtKB |
| cluster19153 | 2 | P43243 | GO:0010608; P:posttranscriptional regulation of gene expression; IDA:CACAO |
| cluster19317 | 2 | Q91VF5 | GO:0010811; P:positive regulation of cell-substrate adhesion; IBA:GO_Central |
| cluster19286 | 2 | Q76HP3 | GO:0010923; P:negative regulation of phosphatase activity; ISS:UniProtKB |
| cluster19206 | 2 | Q9R1J8 | GO:0010976; P:positive regulation of neuron projection development; IDA:ParkinsonsUK-UCL |
| cluster19353 | 2 | Q9ULH0 | GO:0010976; P:positive regulation of neuron projection development; ISS:UniProtKB |
| cluster19365 | 2 | Q13556 | GO:0014733; P:regulation of skeletal muscle adaptation; TAS:UniProtKB |
| cluster18246 | 2 | P70414 | GO:0014829; P:vascular smooth muscle contraction; IMP:BHF-UCL |
| cluster18400 | 2 | P46939 | GO:0014894; P:response to denervation involved in regulation of muscle adaptation; IEA:Ensembl |
| cluster18267 | 2 | A0MGZ7 | GO:0015015; P:heparan sulfate proteoglycan biosynthetic process, enzymatic modification; IBA:GO_Central |
| cluster19301 | 2 | A0MGZ7 | GO:0015015; P:heparan sulfate proteoglycan biosynthetic process, enzymatic modification; IBA:GO_Central |
| cluster18328 | 2 | O60318 | GO:0015031; P:protein transport; IEA:UniProtKB-KW |
| cluster18359 | 2 | Q9NWI9 | GO:0015031; P:protein transport; IEA:UniProtKB-KW |
| cluster18381 | 2 | Q9JIP7 | GO:0015031; P:protein transport; IEA:UniProtKB-KW |
| cluster18406 | 2 | Q811G0 | GO:0015031; P:protein transport; IEA:UniProtKB-KW |
| cluster18465 | 2 | Q9UPQ3 | GO:0015031; P:protein transport; IEA:UniProtKB-KW |
| cluster18508 | 2 | Q9NWI9 | GO:0015031; P:protein transport; IEA:UniProtKB-KW |
| cluster19267 | 2 | O60318 | GO:0015031; P:protein transport; IEA:UniProtKB-KW |
| cluster18293 | 2 | P04323 | GO:0015074; P:DNA integration; IEA:InterPro |
| cluster19221 | 2 | Q96ES6 | GO:0015295; F:solute:proton symporter activity; IBA:GO_Central |
| cluster18374 | 2 | Q0VCM6 | GO:0015804; P:neutral amino acid transport; IBA:GO_Central |
| cluster19316 | 2 | Q06185 | GO:0015986; P:ATP synthesis coupled proton transport; IEA:InterPro |
| cluster18242 | 2 | Q68CR7 | GO:0016021; C:integral component of membrane; IEA:UniProtKB-KW |
| cluster18264 | 2 | Q6ZVL6 | GO:0016021; C:integral component of membrane; IEA:UniProtKB-KW |
| cluster18453 | 2 | Q5GH64 | GO:0016021; C:integral component of membrane; IEA:UniProtKB-KW |
| cluster18474 | 2 | Q6PGS5 | GO:0016021; C:integral component of membrane; IEA:UniProtKB-KW |
| cluster19394 | 2 | Q8BMD6 | GO:0016021; C:integral component of membrane; IEA:UniProtKB-KW |
| cluster18462 | 2 | Q9NPG3 | GO:0016032; P:viral process; IEA:UniProtKB-KW |

| | | | |
|---|---|---|---|
| cluster19315 | 2 | P08047 | GO:0016032; P:viral process; IEA:UniProtKB-KW |
| cluster19293 | 2 | Q80TJ1 | GO:0016050; P:vesicle organization; IMP:MGI |
| cluster19159 | 2 | P43446 | GO:0016055; P:Wnt signaling pathway; IBA:GO_Central |
| cluster19311 | 2 | Q00993 | GO:0016055; P:Wnt signaling pathway; IBA:GO_Central |
| cluster18284 | 2 | B0R0I6 | GO:0016055; P:Wnt signaling pathway; IEA:UniProtKB-KW |
| cluster18336 | 2 | B0R0I6 | GO:0016055; P:Wnt signaling pathway; IEA:UniProtKB-KW |
| cluster19166 | 2 | Q8R4V4 | GO:0016055; P:Wnt signaling pathway; IEA:UniProtKB-KW |
| cluster19230 | 2 | Q6ZSJ9 | GO:0016055; P:Wnt signaling pathway; IEA:UniProtKB-KW |
| cluster19318 | 2 | Q9UKE0 | GO:0016055; P:Wnt signaling pathway; IEA:UniProtKB-KW |
| cluster19347 | 2 | Q9UPM3 | GO:0016055; P:Wnt signaling pathway; IEA:UniProtKB-KW |
| cluster19358 | 2 | Q8QGP3 | GO:0016055; P:Wnt signaling pathway; IEA:UniProtKB-KW |
| cluster18299 | 2 | Q8BY04 | GO:0016082; P:synaptic vesicle priming; IDA:SynGO |
| cluster18537 | 2 | Q62769 | GO:0016082; P:synaptic vesicle priming; IMP:ParkinsonsUK-UCL |
| cluster18263 | 2 | Q1LYM3 | GO:0016192; P:vesicle-mediated transport; IEA:InterPro |
| cluster18471 | 2 | O60229 | GO:0016192; P:vesicle-mediated transport; TAS:ProtInc |
| cluster19340 | 2 | Q9NY28 | GO:0016266; P:O-glycan processing; TAS:Reactome |
| cluster18439 | 2 | Q6GLZ5 | GO:0016476; P:regulation of embryonic cell shape; IMP:UniProtKB |
| cluster18493 | 2 | Q7L273 | GO:0016567; P:protein ubiquitination; IEA:UniProtKB-UniPathway |
| cluster18506 | 2 | Q6PDK8 | GO:0016567; P:protein ubiquitination; IEA:UniProtKB-UniPathway |
| cluster19391 | 2 | Q7L273 | GO:0016567; P:protein ubiquitination; IEA:UniProtKB-UniPathway |
| cluster19410 | 2 | Q96M02 | GO:0016567; P:protein ubiquitination; ISS:UniProtKB |
| cluster18301 | 2 | Q6ZTM0 | GO:0016579; P:protein deubiquitination; IEA:InterPro |
| cluster19275 | 2 | Q6IE24 | GO:0016579; P:protein deubiquitination; IEA:InterPro |
| cluster19403 | 2 | Q91VV4 | GO:0017112; F:Rab guanyl-nucleotide exchange factor activity; ISS:UniProtKB |
| cluster19395 | 2 | A4IGD2 | GO:0017188; F:aspartate N-acetyltransferase activity; IEA:UniProtKB-EC |
| cluster19249 | 2 | Q9JKF6 | GO:0019062; P:virion attachment to host cell; IEA:InterPro |
| cluster18278 | 2 | P62955 | GO:0019226; P:transmission of nerve impulse; IBA:GO_Central |
| cluster18367 | 2 | P62955 | GO:0019226; P:transmission of nerve impulse; IBA:GO_Central |
| cluster19325 | 2 | Q5QT56 | GO:0019227; P:neuronal action potential propagation; IEA:Ensembl |
| cluster18291 | 2 | Q9ET38 | GO:0019227; P:neuronal action potential propagation; IMP:MGI |
| cluster19374 | 2 | Q9NZV8 | GO:0019233; P:sensory perception of pain; IEA:Ensembl |
| cluster18258 | 2 | Q02294 | GO:0019233; P:sensory perception of pain; IMP:RGD |
| cluster18433 | 2 | O55017 | GO:0019233; P:sensory perception of pain; ISO:MGI |

| | | | |
|---|---|---|---|
| cluster19366 | 2 | Q61115 | GO:0021522; P:spinal cord motor neuron differentiation; IGI:MGI |
| cluster18300 | 2 | P23759 | GO:0021527; P:spinal cord association neuron differentiation; IEA:Ensembl |
| cluster18268 | 2 | Q8NFP4 | GO:0021527; P:spinal cord association neuron differentiation; ISS:HGNC |
| cluster18389 | 2 | P70120 | GO:0021537; P:telencephalon development; IGI:MGI |
| cluster19281 | 2 | Q61380 | GO:0021762; P:substantia nigra development; HEP:UniProtKB |
| cluster18295 | 2 | O75093 | GO:0022028; P:tangential migration from the subventricular zone to the olfactory bulb; IEA:Ensembl |
| cluster19341 | 2 | Q8N7C0 | GO:0022414; P:reproductive process; IEA:Ensembl |
| cluster19187 | 2 | A5D7D1 | GO:0030050; P:vesicle transport along actin filament; IEA:Ensembl |
| cluster19188 | 2 | A5D7D1 | GO:0030050; P:vesicle transport along actin filament; IEA:Ensembl |
| cluster18451 | 2 | O43707 | GO:0030050; P:vesicle transport along actin filament; IMP:UniProtKB |
| cluster19387 | 2 | P57780 | GO:0030050; P:vesicle transport along actin filament; ISO:MGI |
| cluster18331 | 2 | Q9R1E6 | GO:0030334; P:regulation of cell migration; ISO:MGI |
| cluster19167 | 2 | Q96FX7 | GO:0030488; P:tRNA methylation; IBA:GO_Central |
| cluster18525 | 2 | Q92859 | GO:0030513; P:positive regulation of BMP signaling pathway; IEA:InterPro |
| cluster19171 | 2 | Q90Y54 | GO:0030878; P:thyroid gland development; IMP:ZFIN |
| cluster18461 | 2 | O95747 | GO:0031098; P:stress-activated protein kinase signaling cascade; IBA:GO_Central |
| cluster18513 | 2 | Q863I2 | GO:0031098; P:stress-activated protein kinase signaling cascade; IBA:GO_Central |
| cluster18413 | 2 | Q62845 | GO:0031175; P:neuron projection development; IDA:UniProtKB |
| cluster19291 | 2 | P22411 | GO:0031295; P:T cell costimulation; ISS:UniProtKB |
| cluster18420 | 2 | Q90YM8 | GO:0031529; P:ruffle organization; ISS:UniProtKB |
| cluster18285 | 2 | O43739 | GO:0032012; P:regulation of ARF protein signal transduction; IEA:InterPro |
| cluster19383 | 2 | O08967 | GO:0032012; P:regulation of ARF protein signal transduction; IEA:InterPro |
| cluster18257 | 2 | Q32NR9 | GO:0032185; P:septin cytoskeleton organization; IMP:UniProtKB |
| cluster18398 | 2 | Q13796 | GO:0032438; P:melanosome organization; ISS:HGNC |
| cluster18372 | 2 | Q96S21 | GO:0032482; P:Rab protein signal transduction; IBA:GO_Central |
| cluster18503 | 2 | Q96S21 | GO:0032482; P:Rab protein signal transduction; IBA:GO_Central |
| cluster19319 | 2 | O08680 | GO:0032496; P:response to lipopolysaccharide; IEP:RGD |
| cluster18469 | 2 | P07812 | GO:0032570; P:response to progesterone; IDA:AgBase |
| cluster18271 | 2 | P15146 | GO:0032570; P:response to progesterone; IEP:RGD |
| cluster19302 | 2 | Q4L1J4 | GO:0032947; F:protein-containing complex scaffold activity; IEA:InterPro |
| cluster19386 | 2 | P34942 | GO:0032981; P:mitochondrial respiratory chain complex I assembly; ISS:UniProtKB |
| cluster18480 | 2 | Q8C8R3 | GO:0033292; P:T-tubule organization; IMP:BHF-UCL |

| | | | |
|---|---|---|---|
| cluster18524 | 2 | P02468 | GO:0034446; P:substrate adhesion-dependent cell spreading; ISO:MGI |
| cluster18294 | 2 | Q7Z3S7 | GO:0034765; P:regulation of ion transmembrane transport; IEA:UniProtKB-KW |
| cluster18394 | 2 | Q99454 | GO:0034765; P:regulation of ion transmembrane transport; IEA:UniProtKB-KW |
| cluster18407 | 2 | Q99244 | GO:0034765; P:regulation of ion transmembrane transport; IEA:UniProtKB-KW |
| cluster18422 | 2 | Q15878 | GO:0034765; P:regulation of ion transmembrane transport; IEA:UniProtKB-KW |
| cluster18463 | 2 | Q99454 | GO:0034765; P:regulation of ion transmembrane transport; IEA:UniProtKB-KW |
| cluster18495 | 2 | Q64347 | GO:0034765; P:regulation of ion transmembrane transport; IEA:UniProtKB-KW |
| cluster19304 | 2 | Q99244 | GO:0034765; P:regulation of ion transmembrane transport; IEA:UniProtKB-KW |
| cluster18438 | 2 | P23735 | GO:0034987; F:immunoglobulin receptor binding; IPI:AgBase |
| cluster18530 | 2 | Q8TEF5 | GO:0035023; P:regulation of Rho protein signal transduction; IEA:InterPro |
| cluster18519 | 2 | Q9Z0U4 | GO:0035094; P:response to nicotine; IEP:RGD |
| cluster18297 | 2 | Q3TYL1 | GO:0035176; P:social behavior; IMP:MGI |
| cluster19224 | 2 | Q3TYL1 | GO:0035176; P:social behavior; IMP:MGI |
| cluster19298 | 2 | Q3TYL1 | GO:0035176; P:social behavior; IMP:MGI |
| cluster18534 | 2 | Q61626 | GO:0035249; P:synaptic transmission, glutamatergic; IMP:MGI |
| cluster19376 | 2 | Q03391 | GO:0035249; P:synaptic transmission, glutamatergic; ISO:MGI |
| cluster18466 | 2 | P42262 | GO:0035249; P:synaptic transmission, glutamatergic; TAS:ARUK-UCL |
| cluster18265 | 2 | Q96MS0 | GO:0035385; P:Roundabout signaling pathway; IEA:InterPro |
| cluster18404 | 2 | Q9JJ00 | GO:0035456; P:response to interferon-beta; ISO:MGI |
| cluster18430 | 2 | O70507 | GO:0035725; P:sodium ion transmembrane transport; ISO:MGI |
| cluster18442 | 2 | Q5XXA6 | GO:0035774; P:positive regulation of insulin secretion involved in cellular response to glucose stimulus; IMP:UniProtKB |
| cluster19284 | 2 | Q6Y636 | GO:0036324; P:vascular endothelial growth factor receptor-2 signaling pathway; IMP:UniProtKB |
| cluster18405 | 2 | Q8CGE9 | GO:0038032; P:termination of G protein-coupled receptor signaling pathway; ISO:MGI |
| cluster18307 | 2 | P56581 | GO:0040029; P:regulation of gene expression, epigenetic; IMP:UniProtKB |
| cluster19369 | 2 | Q9TSZ3 | GO:0042391; P:regulation of membrane potential; IBA:GO_Central |
| cluster18314 | 2 | Q5JR59 | GO:0042803; F:protein homodimerization activity; IPI:UniProtKB |
| cluster19322 | 2 | Q8VC88 | GO:0042803; F:protein homodimerization activity; ISO:MGI |
| cluster18335 | 2 | Q8CG09 | GO:0042908; P:xenobiotic transport; IMP:RGD |
| cluster18396 | 2 | Q9P2T1 | GO:0043101; P:purine-containing compound salvage; TAS:Reactome |
| cluster18440 | 2 | Q9NYB5 | GO:0043252; P:sodium-independent organic anion transport; IBA:GO_Central |
| cluster18489 | 2 | Q9NPD5 | GO:0043252; P:sodium-independent organic anion transport; IBA:GO_Central |

| | | | |
|---|---|---|---|
| cluster18511 | 2 | Q9NPD5 | GO:0043252; P:sodium-independent organic anion transport; IBA:GO_Central |
| cluster18435 | 2 | Q7SY48 | GO:0043487; P:regulation of RNA stability; IMP:ZFIN |
| cluster18429 | 2 | Q63054 | GO:0043496; P:regulation of protein homodimerization activity; IMP:RGD |
| cluster19396 | 2 | Q80U12 | GO:0043547; P:positive regulation of GTPase activity; ISO:MGI |
| cluster18347 | 2 | Q6TEN6 | GO:0043551; P:regulation of phosphatidylinositol 3-kinase activity; ISS:UniProtKB |
| cluster19247 | 2 | Q6TEN6 | GO:0043551; P:regulation of phosphatidylinositol 3-kinase activity; ISS:UniProtKB |
| cluster19308 | 2 | A0ZSK4 | GO:0044179; P:hemolysis in other organism; IEA:UniProtKB-KW |
| cluster18518 | 2 | Q9BSF8 | GO:0044342; P:type B pancreatic cell proliferation; ISS:UniProtKB |
| cluster18486 | 2 | P03360 | GO:0044826; P:viral genome integration into host DNA; IEA:UniProtKB-KW |
| cluster18529 | 2 | Q07954 | GO:0045056; P:transcytosis; IMP:ARUK-UCL |
| cluster19203 | 2 | Q9Y219 | GO:0045061; P:thymic T cell selection; IDA:UniProtKB |
| cluster19404 | 2 | Q6PVZ1 | GO:0045110; P:intermediate filament bundle assembly; ISS:UniProtKB |
| cluster18416 | 2 | Q8CJ99 | GO:0045162; P:clustering of voltage-gated sodium channels; IMP:RGD |
| cluster18542 | 2 | G5E861 | GO:0045162; P:clustering of voltage-gated sodium channels; ISO:MGI |
| cluster18499 | 2 | Q9H604 | GO:0045214; P:sarcomere organization; IEA:Ensembl |
| cluster19236 | 2 | Q9H604 | GO:0045214; P:sarcomere organization; IEA:Ensembl |
| cluster19337 | 2 | Q9H987 | GO:0045214; P:sarcomere organization; ISS:BHF-UCL |
| cluster19170 | 2 | P07221 | GO:0045214; P:sarcomere organization; ISS:UniProtKB |
| cluster18371 | 2 | B6RSP1 | GO:0045218; P:zonula adherens maintenance; ISS:UniProtKB |
| cluster18248 | 2 | P55283 | GO:0045773; P:positive regulation of axon extension; IEA:Ensembl |
| cluster19227 | 2 | P55283 | GO:0045773; P:positive regulation of axon extension; IEA:Ensembl |
| cluster18443 | 2 | Q60821 | GO:0045893; P:positive regulation of transcription, DNA-templated; IDA:MGI |
| cluster19160 | 2 | Q92172 | GO:0045893; P:positive regulation of transcription, DNA-templated; IDA:UniProtKB |
| cluster18515 | 2 | Q08DL5 | GO:0045944; P:positive regulation of transcription by RNA polymerase II; IBA:GO_Central |
| cluster19253 | 2 | O60422 | GO:0045944; P:positive regulation of transcription by RNA polymerase II; IBA:GO_Central |
| cluster19375 | 2 | P17926 | GO:0045944; P:positive regulation of transcription by RNA polymerase II; IBA:GO_Central |
| cluster18476 | 2 | Q03413 | GO:0045944; P:positive regulation of transcription by RNA polymerase II; IEA:InterPro |
| cluster19225 | 2 | Q8NHV1 | GO:0046039; P:GTP metabolic process; IDA:UniProtKB |
| cluster19342 | 2 | P28037 | GO:0046654; P:tetrahydrofolate biosynthetic process; TAS:RGD |
| cluster18290 | 2 | P54760 | GO:0046777; P:protein autophosphorylation; IDA:UniProtKB |

| | | | |
|---|---|---|---|
| cluster18312 | 2 | A5X7A0 | GO:0046872; F:metal ion binding; IEA:UniProtKB-KW |
| cluster18496 | 2 | Q8R0A2 | GO:0046872; F:metal ion binding; IEA:UniProtKB-KW |
| cluster19368 | 2 | A5D6U8 | GO:0046872; F:metal ion binding; IEA:UniProtKB-KW |
| cluster18342 | 2 | Q5I2B1 | GO:0046931; P:pore complex assembly; IEA:InterPro |
| cluster18376 | 2 | O08764 | GO:0046982; F:protein heterodimerization activity; IEA:InterPro |
| cluster18494 | 2 | Q80WQ9 | GO:0046983; F:protein dimerization activity; IEA:InterPro |
| cluster19320 | 2 | Q642M9 | GO:0047115; F:trans-1,2-dihydrobenzene-1,2-diol dehydrogenase activity; IEA:UniProtKB-EC |
| cluster19328 | 2 | Q9HAC7 | GO:0047369; F:succinate-hydroxymethylglutarate CoA-transferase activity; IDA:UniProtKB |
| cluster19272 | 2 | Q9UI94 | GO:0047496; P:vesicle transport along microtubule; IEA:Ensembl |
| cluster18249 | 2 | O14576 | GO:0047496; P:vesicle transport along microtubule; IMP:UniProtKB |
| cluster18363 | 2 | Q5SZD4 | GO:0047961; F:glycine N-acyltransferase activity; IEA:InterPro |
| cluster19229 | 2 | Q5SZD4 | GO:0047961; F:glycine N-acyltransferase activity; IEA:InterPro |
| cluster18279 | 2 | Q8N9V7 | GO:0048137; P:spermatocyte division; IEA:Ensembl |
| cluster18247 | 2 | B8PYG1 | GO:0048168; P:regulation of neuronal synaptic plasticity; IEA:InterPro |
| cluster18272 | 2 | B3DHW5 | GO:0048172; P:regulation of short-term neuronal synaptic plasticity; ISS:UniProtKB |
| cluster19348 | 2 | Q00975 | GO:0048265; P:response to pain; IEA:Ensembl |
| cluster18323 | 2 | Q9WTU6 | GO:0048511; P:rhythmic process; IEA:UniProtKB-KW |
| cluster19209 | 2 | Q91018 | GO:0048511; P:rhythmic process; IEA:UniProtKB-KW |
| cluster19241 | 2 | Q90327 | GO:0048511; P:rhythmic process; IEA:UniProtKB-KW |
| cluster19213 | 2 | P40424 | GO:0048538; P:thymus development; IEA:Ensembl |
| cluster18434 | 2 | A3RK74 | GO:0048745; P:smooth muscle tissue development; IMP:ZFIN |
| cluster19360 | 2 | Q28019 | GO:0050436; F:microfibril binding; IDA:AgBase |
| cluster19279 | 2 | Q9Y2C2 | GO:0050770; P:regulation of axonogenesis; IEA:Ensembl |
| cluster18454 | 2 | P01780 | GO:0050776; P:regulation of immune response; TAS:Reactome |
| cluster18426 | 2 | Q9ET54 | GO:0050808; P:synapse organization; IBA:GO_Central |
| cluster19356 | 2 | Q9ET54 | GO:0050808; P:synapse organization; IBA:GO_Central |
| cluster19321 | 2 | O35927 | GO:0050808; P:synapse organization; ISO:MGI |
| cluster18326 | 2 | Q5PYH5 | GO:0050808; P:synapse organization; NAS:UniProtKB |
| cluster18449 | 2 | Q5PYH5 | GO:0050808; P:synapse organization; NAS:UniProtKB |
| cluster19314 | 2 | Q6R005 | GO:0050808; P:synapse organization; NAS:UniProtKB |
| cluster19389 | 2 | Q5PYH7 | GO:0050808; P:synapse organization; NAS:UniProtKB |

| | | | |
|---|---|---|---|
| cluster18311 | 2 | Q9WU42 | GO:0050872; P:white fat cell differentiation; IMP:MGI |
| cluster18274 | 2 | Q9ESZ0 | GO:0050882; P:voluntary musculoskeletal movement; IEA:Ensembl |
| cluster19393 | 2 | Q9P2F8 | GO:0051056; P:regulation of small GTPase mediated signal transduction; IEA:InterPro |
| cluster19237 | 2 | Q8TF74 | GO:0051127; P:positive regulation of actin nucleation; IBA:GO_Central |
| cluster18478 | 2 | Q9ULH7 | GO:0051145; P:smooth muscle cell differentiation; IBA:GO_Central |
| cluster18308 | 2 | Q1LY46 | GO:0051260; P:protein homooligomerization; ISS:UniProtKB |
| cluster18418 | 2 | Q9IAC6 | GO:0051414; P:response to cortisol; IDA:AgBase |
| cluster19288 | 2 | Q9R1N3 | GO:0051453; P:regulation of intracellular pH; IBA:GO_Central |
| cluster19282 | 2 | Q62920 | GO:0051963; P:regulation of synapse assembly; IDA:UniProtKB |
| cluster19234 | 2 | Q14833 | GO:0051966; P:regulation of synaptic transmission, glutamatergic; IBA:GO_Central |
| cluster19290 | 2 | Q27991 | GO:0055015; P:ventricular cardiac muscle cell development; IEA:Ensembl |
| cluster18349 | 2 | Q5F364 | GO:0055085; P:transmembrane transport; IBA:GO_Central |
| cluster18543 | 2 | Q92887 | GO:0055085; P:transmembrane transport; IBA:GO_Central |
| cluster19292 | 2 | Q8VI47 | GO:0055085; P:transmembrane transport; IBA:GO_Central |
| cluster18340 | 2 | Q0VGW6 | GO:0055085; P:transmembrane transport; IEA:InterPro |
| cluster18288 | 2 | Q9GZV3 | GO:0055085; P:transmembrane transport; TAS:Reactome |
| cluster18483 | 2 | P11615 | GO:0055117; P:regulation of cardiac muscle contraction; IEA:InterPro |
| cluster18380 | 2 | Q9NYQ7 | GO:0060071; P:Wnt signaling pathway, planar cell polarity pathway; NAS:ParkinsonsUK-UCL |
| cluster19248 | 2 | P18845 | GO:0060084; P:synaptic transmission involved in micturition; ISS:UniProtKB |
| cluster19309 | 2 | Q17QN4 | GO:0060307; P:regulation of ventricular cardiac muscle cell membrane repolarization; IEA:Ensembl |
| cluster19367 | 2 | Q9WVQ1 | GO:0060395; P:SMAD protein signal transduction; IDA:UniProtKB |
| cluster18333 | 2 | Q9UDU1 | GO:0060395; P:SMAD protein signal transduction; ISS:UniProtKB |
| cluster18366 | 2 | Q99PP9 | GO:0060416; P:response to growth hormone; ISO:MGI |
| cluster18455 | 2 | P31579 | GO:0060468; P:prevention of polyspermy; IBA:GO_Central |
| cluster18411 | 2 | Q9H9B1 | GO:0060992; P:response to fungicide; IEA:Ensembl |
| cluster18338 | 2 | Q90ZE4 | GO:0061053; P:somite development; IMP:ZFIN |
| cluster18485 | 2 | Q1LVF0 | GO:0061053; P:somite development; IMP:ZFIN |
| cluster18292 | 2 | Q96QZ7 | GO:0065003; P:protein-containing complex assembly; NAS:ProtInc |
| cluster18360 | 2 | A1CQG2 | GO:0070086; P:ubiquitin-dependent endocytosis; IEA:EnsemblFungi |
| cluster18373 | 2 | B0LPN4 | GO:0070296; P:sarcoplasmic reticulum calcium ion transport; IMP:RGD |

| | | | |
|---|---|---|---|
| cluster18488 | 2 | Q9UHV7 | GO:0070328; P:triglyceride homeostasis; IEA:Ensembl |
| cluster19406 | 2 | Q9UHV7 | GO:0070328; P:triglyceride homeostasis; IEA:Ensembl |
| cluster19392 | 2 | O73737 | GO:0070507; P:regulation of microtubule cytoskeleton organization; IBA:GO_Central |
| cluster18479 | 2 | Q86YQ8 | GO:0071277; P:cellular response to calcium ion; IBA:GO_Central |
| cluster19195 | 2 | Q86YQ8 | GO:0071277; P:cellular response to calcium ion; IBA:GO_Central |
| cluster18386 | 2 | O70141 | GO:0071526; P:semaphorin-plexin signaling pathway; IBA:GO_Central |
| cluster18254 | 2 | Q9P2S2 | GO:0071625; P:vocalization behavior; IMP:BHF-UCL |
| cluster19183 | 2 | Q9C081 | GO:0071625; P:vocalization behavior; IMP:BHF-UCL |
| cluster19364 | 2 | Q9P2S2 | GO:0071625; P:vocalization behavior; IMP:BHF-UCL |
| cluster19379 | 2 | Q9P2S2 | GO:0071625; P:vocalization behavior; IMP:BHF-UCL |
| cluster18446 | 2 | Q9CPW0 | GO:0071625; P:vocalization behavior; ISO:MGI |
| cluster19202 | 2 | E9Q7X7 | GO:0071625; P:vocalization behavior; ISO:MGI |
| cluster18415 | 2 | P58392 | GO:0071805; P:potassium ion transmembrane transport; IBA:GO_Central |
| cluster19180 | 2 | Q9UGI6 | GO:0071805; P:potassium ion transmembrane transport; IBA:GO_Central |
| cluster19343 | 2 | Q9UGI6 | GO:0071805; P:potassium ion transmembrane transport; IBA:GO_Central |
| cluster18501 | 2 | Q9Z0U1 | GO:0071847; P:TNFSF11-mediated signaling pathway; IMP:MGI |
| cluster18276 | 2 | P82264 | GO:0072350; P:tricarboxylic acid metabolic process; ISS:UniProtKB |
| cluster18334 | 2 | Q86TV6 | GO:0072659; P:protein localization to plasma membrane; IDA:UniProtKB |
| cluster19215 | 2 | Q8BGB2 | GO:0072659; P:protein localization to plasma membrane; ISS:UniProtKB |
| cluster18390 | 2 | P0C796 | GO:0075732; P:viral penetration into host nucleus; IEA:UniProtKB-KW |
| cluster19175 | 2 | Q92736 | GO:0086005; P:ventricular cardiac muscle cell action potential; ISS:BHF-UCL |
| cluster18330 | 2 | O97831 | GO:0090129; P:positive regulation of synapse maturation; ISS:UniProtKB |
| cluster18516 | 2 | Q80TR1 | GO:0090129; P:positive regulation of synapse maturation; ISS:UniProtKB |
| cluster19333 | 2 | Q3MHM6 | GO:0090136; P:epithelial cell-cell adhesion; IEA:InterPro |
| cluster18325 | 2 | Q91X85 | GO:0097037; P:heme export; IBA:GO_Central |
| cluster18520 | 2 | A4IFW2 | GO:0097374; P:sensory neuron axon guidance; IMP:ZFIN |
| cluster18250 | 2 | Q09YI1 | GO:0098871; C:postsynaptic actin cytoskeleton; IEA:Ensembl |
| cluster19285 | 2 | P70175 | GO:0099072; P:regulation of postsynaptic membrane neurotransmitter receptor levels; IDA:SynGO |
| cluster19163 | 2 | Q8JZP2 | GO:0099504; P:synaptic vesicle cycle; IDA:SynGO |
| cluster19372 | 2 | A5GFW5 | GO:1900026; P:positive regulation of substrate adhesion-dependent cell spreading; ISS:UniProtKB |
| cluster19407 | 2 | P59808 | GO:1900044; P:regulation of protein K63-linked ubiquitination; ISO:MGI |

| | | | |
|---|---|---|---|
| cluster18397 | 2 | Q9Y540 | GO:1900535; P:palmitic acid biosynthetic process; IDA:BHF-UCL |
| cluster18259 | 2 | Q96IZ0 | GO:1901300; P:positive regulation of hydrogen peroxide-mediated programmed cell death; IEA:Ensembl |
| cluster18262 | 2 | Q9QX29 | GO:1902630; P:regulation of membrane hyperpolarization; ISO:MGI |
| cluster18280 | 2 | E9PV86 | GO:1902883; P:negative regulation of response to oxidative stress; ISO:MGI |
| cluster18337 | 2 | P52797 | GO:1902961; P:positive regulation of aspartic-type endopeptidase activity involved in amyloid precursor protein catabolic process; IGI:ARUK-UCL |
| cluster18316 | 2 | Q9HCP3 | GO:1903358; P:regulation of Golgi organization; IDA:UniProtKB |
| cluster18452 | 2 | Q9W6S8 | GO:1904396; P:regulation of neuromuscular junction development; IMP:ZFIN |
| cluster18512 | 2 | G5ECD6 | GO:1904778; P:positive regulation of protein localization to cell cortex; IMP:UniProtKB |
| cluster18531 | 2 | Q9Y2L9 | GO:2000405; P:negative regulation of T cell migration; IMP:UniProtKB |
| cluster18368 | 2 | A5D8S5 | GO:2000564; P:regulation of CD8-positive, alpha-beta T cell proliferation; ISS:UniProtKB |
| cluster18475 | 2 | Q6NRE7 | GO:2000583; P:regulation of platelet-derived growth factor receptor-alpha signaling pathway; ISS:UniProtKB |
| cluster19274 | 2 | Q8IWQ3 | GO:2000807; P:regulation of synaptic vesicle clustering; TAS:ARUK-UCL |
| cluster19307 | 2 | Q6UXZ4 | GO:2001222; P:regulation of neuron migration; IEA:Ensembl |
| cluster18320 | 2 | Q8TB45 | GO:2001236; P:regulation of extrinsic apoptotic signaling pathway; IMP:UniProtKB |
| cluster18507 | 2 | Q9NSC5 | GO:2001256; P:regulation of store-operated calcium entry; IBA:GO_Central |

**Table S2.** Biological process GO terms assigned to the unique clusters identified for *S. tubifer.* The number of assigned to each term is listed.

| ID | Description | Number |
|---|---|---|
| GO:0008150 | biological process | 291 |
| GO:0009987 | cellular process | 178 |
| GO:0065007 | biological regulation | 144 |
| GO:0050896 | response to stimulus | 84 |
| GO:0032501 | multicellular organismal process | 74 |
| GO:0008152 | metabolic process | 71 |
| GO:0032502 | developmental process | 71 |
| GO:0007154 | cell communication | 67 |
| GO:0044237 | cellular metabolic process | 56 |
| GO:0043170 | macromolecule metabolic process | 51 |
| GO:0016043 | cellular component organization | 44 |
| GO:0044238 | primary metabolic process | 42 |
| GO:0006810 | transport | 41 |
| GO:0051234 | establishment of localization | 41 |
| GO:0006807 | nitrogen compound metabolic process | 39 |
| GO:0051179 | localization | 39 |
| GO:0006725 | cellular aromatic compound metabolic process | 34 |
| GO:0046483 | heterocycle metabolic process | 34 |
| GO:0006139 | nucleobase-containing compound metabolic process | 32 |
| GO:0016070 | RNA metabolic process | 25 |
| GO:0051641 | cellular localization | 18 |
| GO:0006464 | cellular protein modification process | 17 |
| GO:0006811 | ion transport | 17 |
| GO:0006928 | movement of cell or subcellular component | 17 |
| GO:0050877 | neurological system process | 17 |
| GO:0051704 | multi-organism process | 17 |
| GO:0006793 | phosphorus metabolic process | 14 |
| GO:0007155 | cell adhesion | 14 |
| GO:0006996 | organelle organization | 13 |
| GO:0015031 | protein transport | 13 |
| GO:0032989 | cellular component morphogenesis | 13 |
| GO:0000003 | reproduction | 9 |
| GO:0002376 | immune system process | 9 |
| GO:0019538 | protein metabolic process | 9 |
| GO:0006259 | DNA metabolic process | 8 |
| GO:0016032 | viral process | 8 |
| GO:0016192 | vesicle-mediated transport | 8 |

| | | |
|---|---|---|
| GO:0040011 | locomotion | 8 |
| GO:0051640 | organelle localization | 8 |
| GO:0006396 | RNA processing | 7 |
| GO:0006508 | proteolysis | 7 |
| GO:0006629 | lipid metabolic process | 7 |
| GO:0065003 | macromolecular complex assembly | 7 |
| GO:0006936 | muscle contraction | 6 |
| GO:0022607 | cellular component assembly | 6 |
| GO:0044255 | cellular lipid metabolic process | 6 |
| GO:0005975 | carbohydrate metabolic process | 5 |
| GO:0008015 | blood circulation | 5 |
| GO:0001775 | cell activation | 4 |
| GO:0009117 | nucleotide metabolic process | 4 |
| GO:0032196 | transposition | 4 |
| GO:0043412 | macromolecule modification | 4 |
| GO:0043603 | cellular amide metabolic process | 4 |
| GO:0044419 | interspecies interaction between organisms | 4 |
| GO:0046903 | secretion | 4 |
| GO:0048511 | rhythmic process | 4 |
| GO:0051186 | cofactor metabolic process | 4 |
| GO:0006082 | organic acid metabolic process | 3 |
| GO:0009116 | nucleoside metabolic process | 3 |
| GO:0015074 | DNA integration | 3 |
| GO:0034622 | cellular macromolecular complex assembly | 3 |
| GO:0040007 | growth | 3 |
| GO:0006260 | DNA replication | 2 |
| GO:0006412 | translation | 2 |
| GO:0006518 | peptide metabolic process | 2 |
| GO:0006836 | neurotransmitter transport | 2 |
| GO:0008283 | cell proliferation | 2 |
| GO:0043500 | muscle adaptation | 2 |
| GO:0046794 | transport of virus | 2 |
| GO:0048469 | cell maturation | 2 |
| GO:0050879 | multicellular organismal movement | 2 |
| GO:0051705 | multi-organism behavior | 2 |
| GO:0001906 | cell killing | 1 |
| GO:0006091 | generation of precursor metabolites and energy | 1 |
| GO:0006281 | DNA repair | 1 |
| GO:0006304 | DNA modification | 1 |
| GO:0006354 | DNA-templated transcription, elongation | 1 |

| | | |
|---|---|---|
| GO:0006805 | xenobiotic metabolic process | 1 |
| GO:0006818 | hydrogen transport | 1 |
| GO:0006865 | amino acid transport | 1 |
| GO:0006914 | autophagy | 1 |
| GO:0007005 | mitochondrion organization | 1 |
| GO:0007030 | Golgi organization | 1 |
| GO:0007049 | cell cycle | 1 |
| GO:0007588 | excretion | 1 |
| GO:0009914 | hormone transport | 1 |
| GO:0015833 | peptide transport | 1 |
| GO:0015849 | organic acid transport | 1 |
| GO:0016049 | cell growth | 1 |
| GO:0016050 | vesicle organization | 1 |
| GO:0022406 | membrane docking | 1 |
| GO:0022610 | biological adhesion | 1 |
| GO:0031640 | killing of cells of other organism | 1 |
| GO:0042180 | cellular ketone metabolic process | 1 |
| GO:0042254 | ribosome biogenesis | 1 |
| GO:0042908 | xenobiotic transport | 1 |
| GO:0043101 | purine-containing compound salvage | 1 |
| GO:0048753 | pigment granule organization | 1 |
| GO:0051181 | cofactor transport | 1 |
| GO:0051258 | protein polymerization | 1 |
| GO:0051276 | chromosome organization | 1 |
| GO:0051301 | cell division | 1 |
| GO:0051703 | intraspecies interaction between organisms | 1 |

**Table S3.** GenBank accession numbers NCBI for phylo tree

| Accession | Species | Family |
|---|---|---|
| MH678615.1 | Acentrogobius caninus | Gobiiformes |
| MG744345.1 | Eugnathogobius polylepis | Gobiiformes |
| MG018480.1 | Gymnogobius petschiliensis | Gobiiformes |
| MK409978.1 | Istigobius campbelli | Gobiiformes |
| MH678617.1 | Paratrypauchen microcephalus | Gobiiformes |
| MF663787.1 | Tridentiger obscurus | Gobiiformes |
| AP005996.1 | Apogon semilineatus | Kurtiformes |
| MH102356.1 | Cheilodipterus quinquelineatus | Kurtiformes |
| MN937193.1 | Jaydia lineata | Kurtiformes |
| AP006030.1 | Kurtus gulliveri | Kurtiformes |
| MN381712.1 | Ostorhinchus fleurieu | Kurtiformes |
| MW007385.1 | Ostorhinchus novemfasciatus | Kurtiformes |
| AP018928.1 | Pristicon trimaculatus | Kurtiformes |
| AP005997.1 | Pterapogon kauderni | Kurtiformes |
| AP018927.1 | Sphaeramia orbicularis | Kurtiformes |
| MF541546.1 | Hippocampus hippocampus | Syngnathiformes |
| MF663787.1 | Trachyrhamphus serratus | Syngnathiformes |
